# Executive control in Obsessive-Compulsive Disorder: A worldwide mega-analysis of task-based functional neuroimaging data of the ENIGMA-OCD consortium

**DOI:** 10.1101/2025.08.20.671231

**Authors:** Nadža Džinalija, Ilya M. Veer, H. Blair Simpson, Iliyan Ivanov, Srinivas Balachander, Francesco Benedetti, Federico Calesella, Sophie M.D.D. Fitzsimmons, Rosa Grützmann, Kristen Hagen, Bjarne Hansen, Stephan Heinzel, Chaim Huijser, Jonathan Ipser, Fern Jaspers-Fayer, Niels T. de Joode, Norbert Kathmann, Minah Kim, Jun Soo Kwon, Wenjuan Liu, Christine Lochner, Ignacio Martinez-Zalacain, Jose M. Menchon, Janardhanan C. Narayanaswamy, Ian S. Olivier, Tjardo S. Postma, Y.C. Janardhan Reddy, Carles Soriano-Mas, S. Evelyn Stewart, Sophia I. Thomopoulos, Anders L. Thorsen, Benedetta Vai, Dick J. Veltman, Ganesan Venkatasubramanian, Valerie Voon, Lea Waller, Ysbrand D. van der Werf, Yi-Jie Zhao, ENIGMA-OCD consortium, Dan J. Stein, Paul M. Thompson, Odile A. van den Heuvel, Chris Vriend

## Abstract

**Objective:** Obsessive-compulsive disorder (OCD) is associated with impaired executive function and altered activity in associated neural circuits, contributing to reduced goal-directed behavior. To investigate neural activation during executive control, we conducted a mega-analysis in the ENIGMA-OCD consortium pooling individual participant data from 475 individuals with OCD and 345 healthy controls across 15 fMRI tasks collected worldwide.

**Methods:** Individual participant data was uniformly processed using HALFpipe to construct voxelwise statistical images of executive control and task load contrasts. Parameter estimates extracted from regions of interest were entered into multilevel Bayesian models to examine regional and whole-brain effects of diagnosis, and, within OCD, the influence of medication status, symptom severity, and age of onset on task activation.

**Results:** We observed a robust task activation pattern across individuals with OCD and control participants in executive control regions across tasks. Relative to controls, individuals with OCD showed moderate to very strong evidence of weaker activation of the dorsolateral prefrontal cortex, precuneus, frontal eye fields, and inferior parietal lobule during executive control (all positive posterior probabilities [P+]<0.1). Individuals with OCD also showed stronger activation in regions of the default mode network during executive function relative to controls. We found little evidence for differential activation during executive control in task-positive regions related to disease onset, severity and medication status.

**Conclusion:** In the first mega-analysis of fMRI studies of executive function in OCD, we found strong evidence of weaker frontoparietal activation during executive control tasks. Our findings also suggest a failure of default mode network regions to appropriately disengage during task performance in OCD.

## Introduction

Obsessive-compulsive disorder (OCD) involves persistent, distressing thoughts (obsessions) and repetitive behaviors or mental acts (compulsions) performed to reduce distress or prevent feared outcomes. These behaviors are thought to arise partly from impairments in executive control, a combination of sub-constructs of the *cognitive systems* domain of the Research Domain Criteria (RDoC(1)). Executive control enables goal-directed behavior and depends on higher-order cognitive functions such as working memory (to hold and manipulate information) and planning (to think and act on that information). Executive dysfunction has been suggested as a possible endophenotype of OCD, representing a heritable mediator trait between genotype and phenotype that acts as a risk factor for the disorder(2, 3).

Tower of London (TOL) and n-back tasks are often used during MRI to study the neural correlates of executive control. These tasks primarily engage the dorsal cognitive circuit, including the dorsolateral prefrontal cortex (dlPFC), pre-supplementary motor area (pre-SMA), and dorsal caudate(4). During these tasks, top-down regulatory control of behavior is provided by the frontoparietal network, encompassing the dlPFC, posterior parietal cortex, anterior cingulate, and insula, through connections with other networks including the dorsal cognitive circuit(5). Individuals with OCD typically perform worse on these tasks, with small-to-medium effect sizes for behavioral performance(5–7), while neuroimaging findings are less consistent. Meta-analyses have found less task-related activation in individuals with OCD than in controls in the frontal and cingulate gyrus, caudate(8–10), precuneus(8), postcentral and inferior occipital gyrus(9), and putamen(10) during executive function. However, stronger activation has also been reported in OCD during executive tasks in the posterior insula/putamen, precentral gyrus, dlPFC, and cuneus(9). As these meta-analyses combined executive and inhibitory control paradigms (where an automatic or prepotent motor response must be withheld), the cognitive construct most linked to OCD-related alterations remains unclear.

Because executive function and inhibitory control rely on partially distinct fronto-striatal networks(4), and executive control encompasses a broader spectrum of goal-directed cognitive processes, combining these paradigms may have obscured their specific neural underpinnings. Isolating executive control tasks is therefore crucial to advance our understanding of impaired goal-directedness in OCD independent of inhibitory control deficits.

The Enhancing Neuro-Imaging and Genetics through Meta-Analysis (ENIGMA) consortium’s OCD Working Group pools participant-level neuroimaging data into large, clinically and ethnically diverse datasets that allow well-powered studies(11). We recently demonstrated for the first time in psychiatry a proof-of-concept that individual-participant fMRI data from different task paradigms within another RDoC domain, i.e., negative valence, can be combined into a single large group-level analysis to compare domain-relevant activation between cases and controls and to investigate clinical subgroups of OCD(12). This subject-level approach to mega-analysis using whole-brain statistical maps provides a richer and more accurate characterization of activation than traditional coordinate-based meta-analyses(13, 14). Given the available task data, the ENIGMA-OCD consortium is uniquely positioned to investigate the neural correlates of executive function separately from inhibitory control.

Here, we isolated the executive control paradigms, including visuo-spatial planning, working memory, and task-switching, in order to study brain circuit function during executive control in OCD independent from inhibitory control. We pooled individual participant data from 15 different global samples that acquired fMRI data during executive control tasks in individuals with OCD and healthy controls (both adults and children) into one image-based mega-analysis. We further investigated associations with relevant clinical characteristics of OCD, such as symptom severity, age of OCD onset, and medication status. Serotonin reuptake inhibitors (SRIs) were previously shown to normalize dorsal attention and default-mode network (DMN) activation during a TOL task in adults with OCD(15). Earlier onset of OCD and greater symptom severity were correlated with worse performance on a visuo-spatial memory task(16) and Tower of Hanoi task(17), respectively. We hypothesized that relative to healthy controls, individuals with OCD would show weaker activation during executive tasks in brain regions related to executive control, and that this difference would increase with increasing task load. We expected that weaker executive control-related activation would be associated with an unmedicated status, early onset of OCD, and greater symptom severity.

## Methods

### Study population

We included 15 samples (seven unpublished), from the ENIGMA-OCD consortium, an international collaboration between sites with legacy historical neuroimaging data from individuals with OCD as well as healthy controls (HCs). The samples spanned ten countries across four continents. Trained personnel administered diagnostic tools based on DSM criteria to diagnose OCD, while OCD severity was measured using the (Children’s) Yale-Brown Obsessive-Compulsive Scale ((C)Y-BOCS;(18, 19)). HCs had no psychiatric diagnosis or psychotropic medication use. Each sample applied its own inclusion and exclusion criteria based on the original study’s research questions and task performance cutoffs; we maintained these criteria in the current analysis. At each participating site, participants provided informed consent, and local review boards approved the use of de-identified data by the ENIGMA-OCD consortium.

### Executive control tasks

Participants in each sample completed one of the following executive control tasks in the MRI scanner: Tower of London (7 samples), n-back (5 samples), and set-shifting (3 samples) tasks (Table 1). Each task included trials testing executive control, enabling us to create one common contrast across paradigms: cognitive load versus no load. We previously used a similar approach for negative valence paradigms(12). Many of these tasks (9 samples) additionally included different difficulty levels, enabling us to also create a contrast that models increased cognitive effort with increasing task load.

**Table 1.**
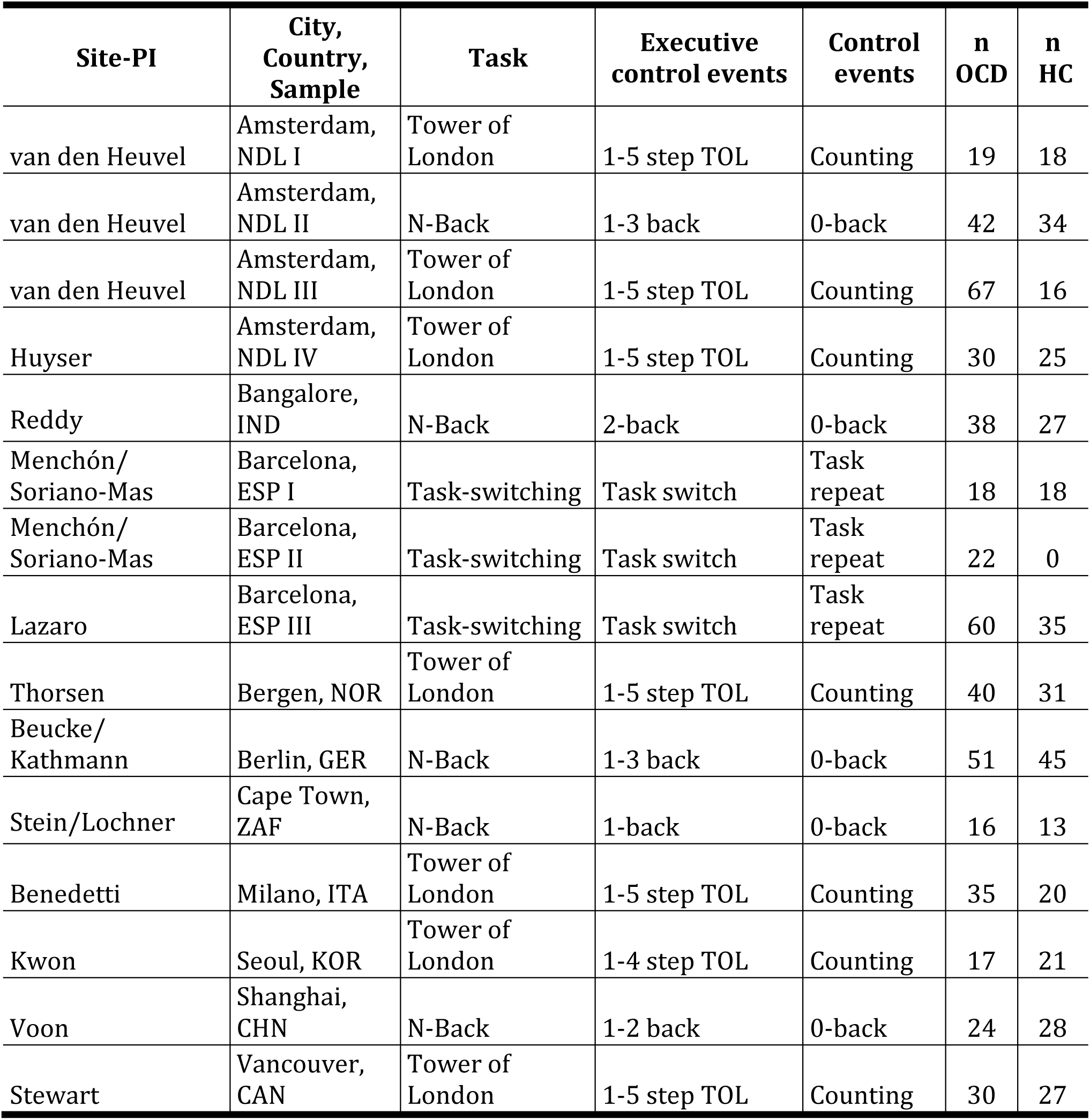
Task characteristics of studies included in mega-analysis.

| **Site-PI** | **City,**<br>**Country,**<br>**Sample** | **Task** | **Executive**<br>**control events** | **Control**<br>**events** | **n**<br>**OCD** | **n**<br>**HC** |
| --- | --- | --- | --- | --- | --- | --- |
| van den Heuvel | Amsterdam,<br>NDL I | Tower of<br>London | 1-5 step TOL | Counting | 19 | 18 |
| van den Heuvel | Amsterdam,<br>NDL II | N-Back | 1-3 back | 0-back | 42 | 34 |
| van den Heuvel | Amsterdam,<br>NDL III | Tower of<br>London | 1-5 step TOL | Counting | 67 | 16 |
| Huyser | Amsterdam,<br>NDL IV | Tower of<br>London | 1-5 step TOL | Counting | 30 | 25 |
| Reddy | Bangalore,<br>IND | N-Back | 2-back | 0-back | 38 | 27 |
| Menchón/<br>Soriano-Mas | Barcelona,<br>ESP I | Task-switching | Task switch | Task<br>repeat | 18 | 18 |
| Menchón/<br>Soriano-Mas | Barcelona,<br>ESP II | Task-switching | Task switch | Task<br>repeat | 22 | 0 |
| Lazaro | Barcelona,<br>ESP III | Task-switching | Task switch | Task<br>repeat | 60 | 35 |
| Thorsen | Bergen, NOR | Tower of<br>London | 1-5 step TOL | Counting | 40 | 31 |
| Beucke/<br>Kathmann | Berlin, GER | N-Back | 1-3 back | 0-back | 51 | 45 |
| Stein/Lochner | Cape Town,<br>ZAF | N-Back | 1-back | 0-back | 16 | 13 |
| Benedetti | Milano, ITA | Tower of<br>London | 1-5 step TOL | Counting | 35 | 20 |
| Kwon | Seoul, KOR | Tower of<br>London | 1-4 step TOL | Counting | 17 | 21 |
| Voon | Shanghai,<br>CHN | N-Back | 1-2 back | 0-back | 24 | 28 |
| Stewart | Vancouver,<br>CAN | Tower of<br>London | 1-5 step TOL | Counting | 30 | 27 |

Due to task design variability, performance metrics were not directly comparable across tasks. Therefore, we restricted our analysis of task performance to the TOL tasks, where sufficient consistency allowed for meaningful comparisons (see Supplemental Methods).

### MRI Image acquisition and processing

MRI data acquisition was not prospectively harmonized; sites contributed pre-existing data (Table S1). Instead, we harmonized image processing across samples using the open-source containerized Harmonized AnaLysis of Functional MRI pipeline (HALFpipe;(20)) version 1.2.2 (see Supplemental Methods) to preprocess the structural and functional images and extracting the task contrasts of interest. HALFpipe was specifically designed to accommodate multi-site fMRI data with heterogeneous acquisition parameters, including differences in field strength and voxel size. Using default settings within HALFpipe, structural pre-processing included skull stripping, tissue segmentation, and spatial normalization while functional pre-processing included motion correction and calculation of 6 rigid-body motion parameters, slice time correction (if slice acquisition order was known), susceptibility distortion correction (if fieldmaps were available), coregistration, spatial normalization, smoothing with a 6mm FWHM Gaussian kernel and extraction of noise components with Independent Component Analysis-based Automatic Removal of Motion Artifacts (ICA-AROMA;(21)).

We excluded participants with mean framewise displacement >1.0 mm. Processed data was also subjected to a quality assessment at each site according to harmonized guidelines (see Supplemental Methods).

### First-level contrast parameter estimate maps

The first-level contrast of interest across tasks compared conditions that test executive functions (planning trials in the TOL task, working memory trials in the n-back task, and switch trials in task-switching task) with baseline conditions (counting trails in the TOL task, 0-back trials in the n-back task, and non-switch trails in task-switching task) (Table S2). The task load contrast modeled increasing cognitive effort, capturing step-wise variations in cognitive demand across conditions.

### Analyses

The analysis plan and hypotheses were pre-registered at <u>osf.io/ebtpk</u> and analysis scripts are available at <u>github.com/nadza-dz/task-based-fMRI-processing-pipeline-ENIGMA-OCD.git</u>. We used the same processing and analysis pipeline as in Dzinalija et al. (12) on negative valence processing.

### ROI analyses

We investigated the regions of interest (ROI) identified in Nitschke et al.’s(22) meta-analysis of the TOL task in HCs (including both adult and pediatric samples): dorsolateral prefrontal cortex (dlPFC), frontal eye fields (FEF), (pre-)supplementary motor area ((pre-)SMA), inferior parietal lobule (IPL), precuneus, caudate, anterior insula, rostrolateral prefrontal cortex (rlPFC), posterior cingulate, inferior occipital gyrus (IOG), and inferior temporal gyrus (ITG) (Table S3). These ROIs were selected based on their involvement in the TOL task; however, they are also relevant to other executive control tasks, as they form part of common neural networks underlying cognitive control, planning, and problem-solving(23). The coordinates of peak activation in cortical regions identified by Nitschke et al. were warped to MNI152 NLIN 2009c (asymmetric) space and a sphere of 5 mm was created around each coordinate. The Melbourne subcortical atlas scale 1(24) was used to identify the caudate. The mean activation of all voxels within an ROI was extracted from the z-statistic maps of each participant’s first-level executive and task load contrasts, provided there was signal in >30% of voxels in the ROI (as done in(12)). For ROIs for which two MNI coordinates exist per hemisphere, we created spheres at each coordinate and averaged activation across all voxels within the two spheres.

Rather than applying separate linear models to each ROI, all regions were entered simultaneously into one Bayesian Multilevel model (RBA, v1.0.10;(25)) that considers the shared non-independent information across brain regions (derived from one participant) in one statistical model. Using a noninformative Gaussian prior, the model was estimated with four Markov chains of 4,000 iterations each, and convergence was confirmed by Rhat < 1.1. A detailed discussion of the advantages of this approach is provided in Dzinalija et al.(12) and Chen et al.(25), but briefly, this approach captures the complex dependencies in the data, dissolves the multiple testing problem, and – unlike null hypothesis significance testing – allows us to test the credibility of our hypotheses directly. We infer the credibility of evidence of an effect from the area under the positive posterior distribution (see Supplemental Methods), summarized as the positive posterior probability (P+), with values <0.10 or >0.90 indicating moderate evidence, <0.05 or >0.95 indicating strong evidence, and <0.025 or >0.975 indicating very strong evidence of an effect(25).

We investigated the main effect of executive function across all tasks and participants (OCD and HCs). Our main effect of interest was the case-control effect, with further analyses for clinical subgroups of OCD. To investigate the effect of age at OCD onset, we performed pairwise comparisons of early-onset OCD individuals (age of onset<18), late-onset OCD individuals (age of onset≥18), and HCs, in adults only. For all other analyses we grouped adults (age≥18) and children (age<18). We investigated the effect of current medication status by pairwise comparisons of medicated individuals (taking any psychotropic medication), unmedicated individuals, and HCs. We assessed the effect of symptom severity using (C)Y-BOCS score as a continuous variable. Sample, age, and sex were entered as covariates of no interest in each analysis.

### Sensitivity analyses

This study is not intended as a methods comparison, therefore we performed select sensitivity analyses specifically to assess the robustness of our findings in line with the primary aim of investigating executive control in OCD. For each ROI analysis on the executive contrast, we performed a leave-one-sample-out analysis to test the robustness of our results to sample effects. We also compared Bayesian multilevel results for the executive contrast with a classical frequentist inference approach using a comparable multilevel mixed-effects linear model with sample and participant as random effects. To assess the influence of sample correction method, we compared Bayesian results that incorporate sample as a covariate of no interest with those obtained using ComBat harmonization for the executive contrast of case-control effects.

Finally, we considered potential task effects in two ways. First, we selectively examined the contrasts of interest within the TOL tasks to determine whether the observed activation patterns were consistent with previous literature, as TOL tasks are most commonly used to assess executive control. Second, we explored including task as a covariate of no interest to account for potential task-related variance across contrasts. Including both task and sample as covariates was not feasible due to near-perfect collinearity between sample and task, which prevented model convergence. Therefore, we included only task as a covariate in this sensitivity analysis.

### Whole-brain analyses

In order to apply the Bayesian multilevel model to the entire brain, whole-brain analyses were done in functionally-defined anatomical parcellations based on the Schaefer-Yeo 7-network 200-parcel atlas(26) and the Melbourne subcortical atlas scale 2(24). Z-statistic maps of each participant’s first-level contrasts were parcellated and activation was entered into identical Bayesian multilevel group analyses as explained above. To determine the main effect of executive function across all tasks, we used an intercept model pooling all participants (HCs and individuals with OCD) during executive control. Covariates of no interest were again sample, sex and age.

## Results

### Participants

The final sample included 475 participants with OCD (51.2% female, mean age=29.4±12.4) and 345 HCs (52.8% female, mean age=28.7±11.9)) (Table 2, S4). See Supplemental Methods for exclusions (n=16 OCD, n=25 HCs) and sample sizes per analysis.

**Table 2.** Demographics of full sample. Data are expressed as mean (SD) unless otherwise noted. Medication status is measured at time of scan. OCD = obsessive-compulsive disorder; HC = healthy control; (C)Y-BOCS = (Children’s) Yale-Brown Obsessive-Compulsive Scale.

|  | OCD<br>n=475 | HC<br>n=345 | Statistic |
| --- | --- | --- | --- |
| **Sex** |  |  |  |
| n female (%) | 243 (51.16%) | 182 (52.75%) |  |
| n male (%) | 232 (48.84%) | 163 (47.25%) | $X^2_{(1)} = 0.14, P = 0.7$ |
| **Age** | 29.4 (12.42) | 28.7 (11.91) |  |
| n <18 (%) | 101 (21.26%) | 67 (19.42%) |  |
| n ≥18 (%) | 374 (78.74%) | 278 (80.58%) | $T_{(818)} = -0.81, P = 0.42$ |
| **Age of OCD onset** |  |  |  |
| n onset <18 (%) | 311 (65.47%) | - |  |
| n onset ≥18 (%) | 143 (30.11%) | - |  |
| n missing data (%) | 17 (3.58%) | - |  |
| **Medication** |  |  |  |
| n medicated (%) | 234 (49.26%) | - |  |
| n unmedicated (%) | 240 (50.53%) | - |  |
| n missing data (%) | 1 (0.21%) | - |  |
| **(C)Y-BOCS** | 23.73 (7.29) | - |  |
| n missing data (%) | 6 (1.26%) | - |  |

### Task Performance

Accuracy was comparable between groups in TOL tasks, except in one sample where individuals with OCD had lower performance than HCs (Table S5).

### Task effects: Executive control and task load

We compared activation during executive control relative to the baseline condition across all participants (cases and controls), both for all tasks combined and specifically for TOL tasks (Figure 1A). Across all tasks, there was very strong evidence of stronger activation in all predefined ROIs during executive control (all P+≥0.99) except the right inferior occipital gyrus (IOG; P+=0.53) and left cingulate gyrus (P+<0.001). Within TOL tasks, there was very strong evidence of weaker activation in the bilateral IOG (P+<0.001) and stronger activation in all other ROIs (P+>0.99). There was stronger task-load-related activation in all ROIs (Figure S1A), both in TOL tasks and all tasks together (P+>0.99).

**Figure 1.**
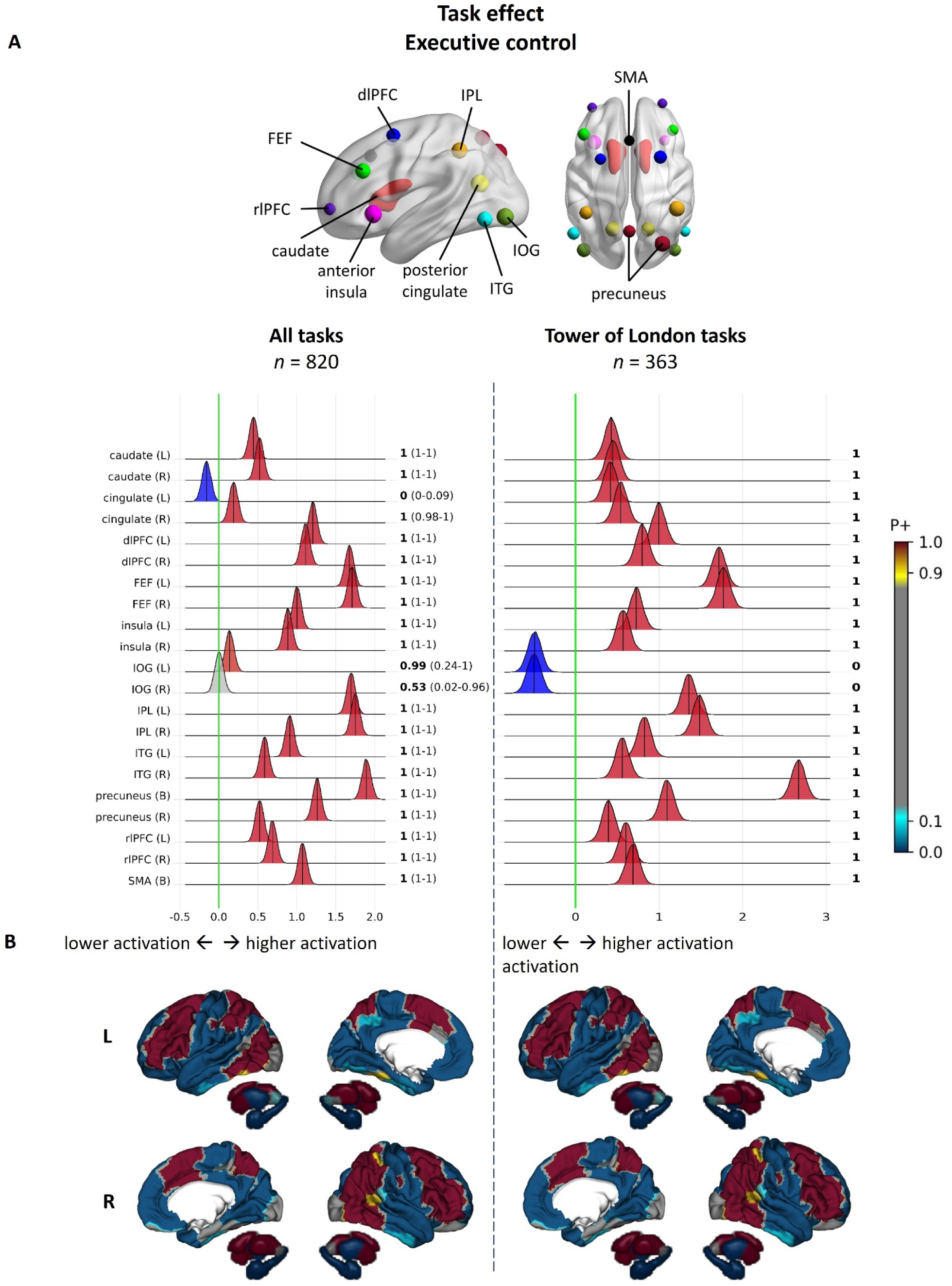
Main effect of executive control across all participants. (A) Region-of-interest results from Bayesian multilevel analyses of group-level executive contrast across individuals with OCD and HCs in all tasks (N tasks = 15, n participants = 820) and in only Tower of London tasks (N tasks = 7, n participants = 363). Posterior probability distributions express the credibility of an effect in each region. Next to each distribution the posterior probability of a positive effect (P+) is shown in bold, as well as the range of values this probability took on in leave-one-sample-out sensitivity analyses (where this analysis was carried out). In regions with posterior distributions to the right of the green no-effect line there is evidence of stronger activation during executive control while in regions with posterior distributions to the left of this line there is evidence of weaker activation during executive control. Regions are color- coded to reflect the strength of evidence for an effect, where in (darker) red regions there is stronger evidence of activation during executive control (P+ values >0.90 indicate moderate to very strong evidence for a positive effect). In (darker) blue regions there is stronger evidence of deactivation during executive control (P+ values <0.10 indicate moderate to very strong evidence for a negative effect). In grey regions there is no strong evidence of activation or deactivation during executive control. Values on the x-axis represent the difference in regional activation levels between executive and control trials (expressed as difference in Z-scores). (B) Whole-brain effects (P+ values) from Bayesian multilevel analyses. P+ values derived from Bayesian multilevel analyses denote the probability that there is stronger brain activation in a given region of the Schaefer 200-parcel 7-network cortical atlas and Melbourne 32-region subcortical atlas. Displayed are lateral and medial views of the cortex. dlPFC= dorsolateral prefrontal cortex; FEF = frontal eye fields; IOG = inferior anterior occipital gyrus; IPL = inferior parietal lobule; ITG = inferior temporal gyrus; rlPFC = rostrolateral prefrontal cortex; SMA = supplementary motor area.

Whole-brain results confirmed task-positive activation in frontoparietal and dorsal attention regions, and showed weaker activation in prefrontal and temporal regions of the DMN for both contrasts (Figure 1B,S1B).

### Brain activity during executive function in OCD versus HCs

In ROI analyses, individuals with OCD, relative to HCs, showed moderate evidence of weaker activation in bilateral dlPFC, left FEF, left ITL, right precuneus, and bilateral (pre-)SMA (0.05<P+≤0.1), and strong-to-very-strong evidence of weaker activation in right FEF, bilateral IPL, and bilateral precuneus (0.01<P+<0.03) during executive control in all tasks (Figure 2A). Results were very similar when considering only TOL tasks. In the task load contrast, there was moderate evidence for weaker activation in individuals with OCD relative to HCs in bilateral caudate, right dlPFC, bilateral FEF, left IPL, bilateral precuneus, and SMA (0.06<P+<0.09) when investigating all tasks, but not when looking at TOL tasks alone (Figure S2A).

**Figure 2.**
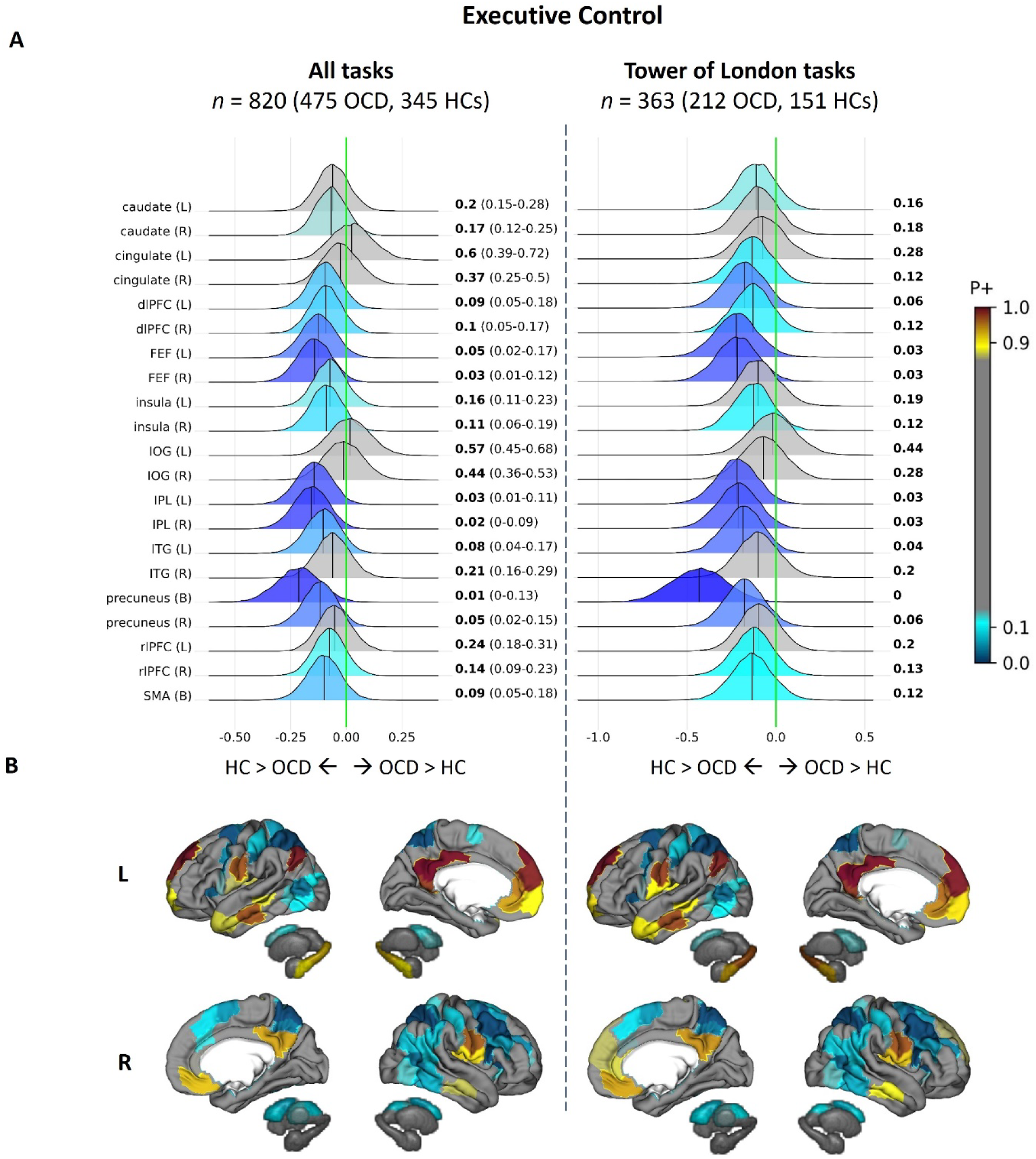
Case-control differences in executive control. (A) Region-of-interest results from Bayesian multilevel analyses. Distributions to the right of the green no-effect line represent regions in which individuals with OCD show evidence for stronger activation than HCs. (Darker) red color represents regions in which individuals with OCD show moderate-to-very- strong evidence for stronger activation than HCs. Regions with posterior distributions to the left of the no-effect line show evidence for stronger activity in HCs than in OCD. (Darker) blue color represents regions in which HCs show moderate-to-very-strong evidence for stronger activation than OCD individuals. In gray-colored regions there is no evidence of a difference between HCs and OCD individuals. Values on the x-axis represent the difference in regional activation levels between HCs and OCD (expressed as difference in Z-scores). (B) Whole-brain effects from Bayesian multilevel analyses. dlPFC= dorsolateral prefrontal cortex; FEF = frontal eye fields; IOG = inferior anterior occipital gyrus; IPL = inferior parietal lobule; ITG = inferior temporal gyrus; rlPFC = rostrolateral prefrontal cortex; SMA = supplementary motor area.

In whole-brain analyses, both the executive and task load contrasts showed evidence for weaker activation in dorsal attention and frontoparietal regions in OCD. The executive contrast additionally showed evidence of stronger activation in default mode regions (i.e., medial prefrontal and posterior cingulate, see Figure 2B,S2B).

### Executive function in clinical subgroups of OCD – age of onset, medication status, and symptom severity effects

#### Age of OCD onset

There was moderate evidence of weaker activation during executive control in adults with early-onset OCD than in adults with late-onset OCD in the bilateral precuneus (P+=0.90) (Figure 3A), but not in any other pre-defined ROIs. Whole-brain analyses demonstrated widespread weaker activation in early-onset OCD across prefrontal, parietal, and temporal regions relative to late-onset OCD. These effects were absent for task load-related activation.

**Figure 3.**
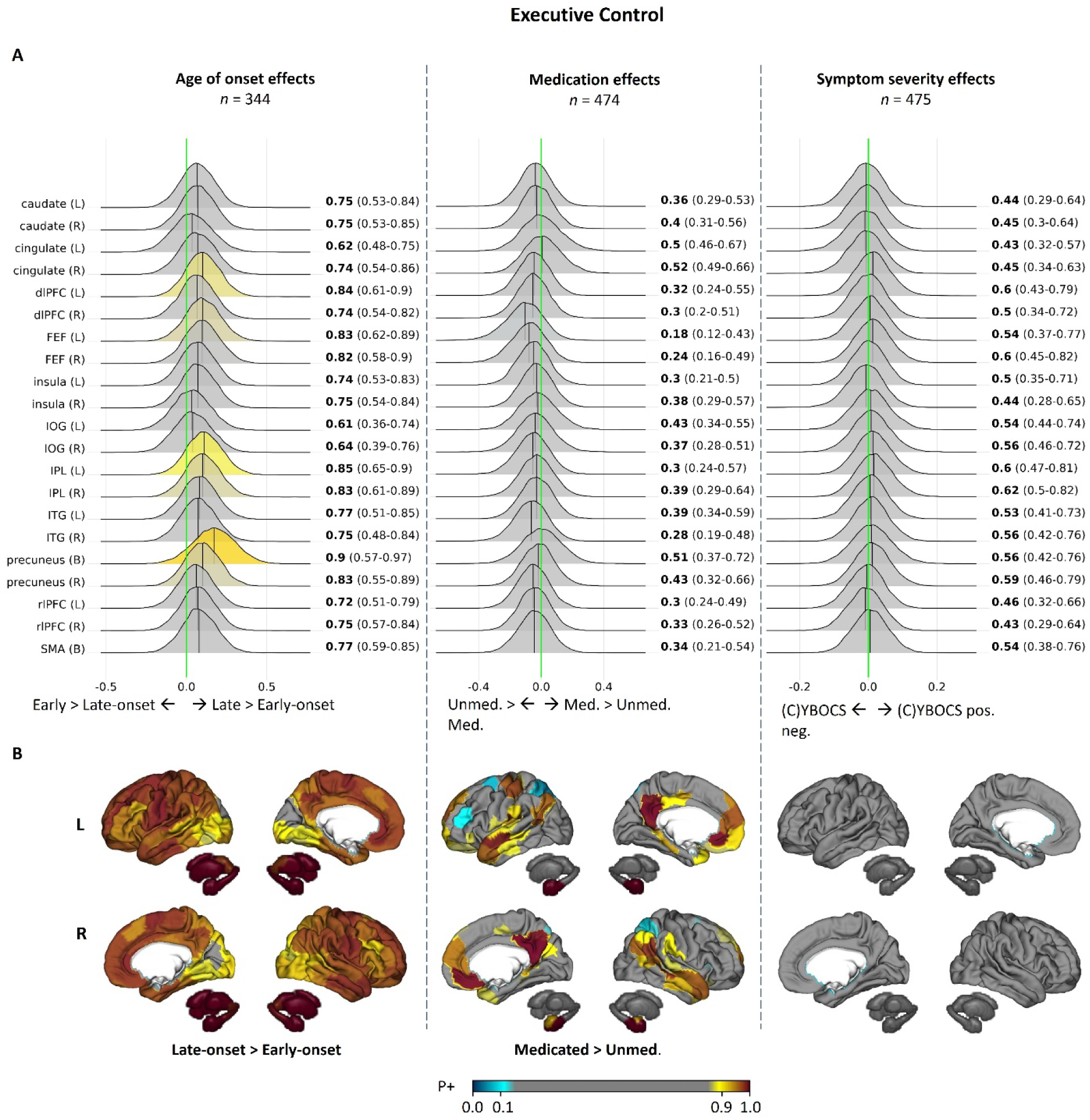
Effects of clinical features of OCD on executive control. (A) Region-of-interest effects of age of onset, medication status, and symptom severity during executive control. (B) Whole-brain analyses of executive control. (C)YBOCS = (Children’s) Yale-Brown Obsessive- Compulsive Scale; dlPFC= dorsolateral prefrontal cortex; FEF = frontal eye fields; IOG = inferior anterior occipital gyrus; IPL = inferior parietal lobule; ITG = inferior temporal gyrus; rlPFC = rostrolateral prefrontal cortex; SMA = supplementary motor area.

#### Medication status

There was no evidence for different activation in any pre-defined ROIs during executive control between medicated and unmedicated individuals with OCD (Figure 3). Whole-brain analyses showed moderate evidence for weaker activation in orbitofrontal, posterior cingulate, and temporal pole regions during executive control in unmedicated than medicated individuals. None of these results were found for task-load related activation, either in ROI or whole-brain analyses.

#### Symptom severity

There was no evidence of an association between symptom severity and brain activation during executive control, either in ROI or whole-brain analyses (Figure 3). There was, however, moderate to very-strong evidence that greater symptom severity correlated with stronger task load-related activation in all investigated ROIs (all P+>0.90), except the left insula (Figure S3A). Whole-brain analyses indicated that OCD severity was associated with stronger activation in dorsal prefrontal and inferior parietal regions (Figure S3B).

### Sensitivity analyses

Leave-one-sample-out validation showed low variability in case-control analyses and clinical subgroup analyses, indicating high robustness of results to sample-specific effects (Figures S5). Results were extremely similar in frequentist statistical models (Table S6) as in Bayesian models. Sample correction using ComBat yielded results nearly identical to those obtained by modeling sample as a covariate (Figure S6). Similarly, including task as a covariate instead of sample yielded results nearly identical to the original model, indicating that case-control effects were robust to potential task-related variance (Figure S6).

## Discussion

Using a mega-analytic approach, we examined neural differences in the RDoC *cognitive systems* domain between OCD and HCs across 15 global samples using participants’ whole-brain executive control- and task load-related activation maps. In almost all ROIs relevant for executive control, we found strong task activation, indicating that we successfully captured domain-specific activation during executive control. In line with our hypotheses, we found weaker activation in OCD in prefrontal, inferior temporal, precuneus, inferior parietal, frontal eye field, and (pre-)supplementary motor areas. With increasing task load, individuals with OCD showed weaker activation than HCs in these regions, and in the caudate and dorsolateral prefrontal cortex. Contrary to expectations and our previous findings on negative valence processing(12), clinical heterogeneity in OCD had little impact on activation strength during executive tasks, though we found some evidence that early-onset OCD may be associated with weaker executive control-related activation, while greater symptom severity appears linked to stronger task load activation.

Weaker activation in task-relevant brain regions in individuals with OCD relative to controls was reported by multiple meta-analyses(8–10), specifically in the frontal gyri and precuneus, closely matching our results. These regions are considered part of the frontoparietal and dorsal cognitive networks(4) and are involved in goal-directed behavior, possibly relying on the precuneus, inferior parietal lobule and frontal eye fields to guide attention processes(27–29) and on the dorsolateral prefrontal cortex and inferior temporal gyrus to integrate multimodal sensory information to guide decision-making(30, 31). This supports the hypothesis that decreased activation during executive control in OCD mediates the inability of individuals with OCD to override compulsions and act in a goal-directed way.

Though clinical features (medication status, age of onset, and symptom severity) showed no strong associations with executive control-related activation in predefined ROIs, whole-brain analyses revealed weaker activation in frontoparietal and temporal regions in early-onset OCD compared to late-onset OCD. This aligns with our hypothesis and earlier findings(16) that early-onset OCD is associated with weaker executive control activation, though it partly conflicts with meta-analytic findings that individuals with early-onset OCD perform worse on visual memory tasks but not on verbal memory or executive control tasks outside the scanner(32). Due to task design heterogeneity, we could not investigate performance differences between OCD and HCs across all tasks, nor relate observed brain activation differences to performance. In TOL tasks, only one sample showed accuracy differences between OCD and HCs, suggesting that activation differences are unlikely to be driven by accuracy, though we cannot rule out differences in other performance metrics such as response times.

We found no association between symptom severity and executive control activation, though severity did correlate positively with task load activation. Paradoxically, individuals with more severe OCD showed stronger task load-related activation in dorsal prefrontal and inferior parietal regions, where OCD individuals on average demonstrated weaker activation during executive control relative to HCs. In post-hoc analyses we found a significant positive correlation between older age at OCD onset and greater symptom severity in the sub-sample that was used to study task load activation, possibly indicating that individuals with late-onset OCD were driving the OCD severity effects on task load activation (Figure S7). Interestingly, previous studies did not find an association between OCD severity and brain activation during executive tasks(33), except in the most severe cases (Y-BOCS>32;(34)), further underscoring the idea that phenotypic heterogeneity in OCD also extends to neural activation.

Alongside weaker executive control-related activation, individuals with OCD showed less deactivation in default mode network (DMN) regions, suggesting difficulty in suppressing the DMN and engaging task-relevant networks. While it remains unclear if switching between executive and default processing modes is impaired in OCD, limited research suggests altered DMN activity and connectivity, both at rest and during task performance(35–37). Specifically, prior ENIGMA-OCD analyses revealed weaker resting state functional connectivity within regions of the DMN – specifically between precuneus/posterior cingulate cortex and dorsomedial prefrontal cortex – and between the DMN and other task-relevant networks like the frontoparietal and dorsal attention network(35). Task-based findings align with this pattern, showing reduced DMN connectivity during a reward task (between the posterior cingulate cortex and the ventromedial prefrontal cortex)(36) and less deactivation in frontal DMN regions during an emotional processing task when shifting from rest to viewing pleasant stimuli(37). This suggests that altered within- and between-network connectivity of the DMN may underlie difficulties in suppressing DMN activity to support goal-directed processing, possibly by impairing attentional control(38). A deficit in deactivating the DMN during task execution may not be specific to OCD, as a suppression deficit was also found during a working memory task in participants with remitted major depressive disorder and was related to rumination(39). This supports the theory that self-referential processing and internal regulation states that are governed by the DMN(40) may be disrupted in OCD in a way that impairs the ability to override internally-generated obsessions and resulting compulsions to act in a goal-directed way. Further research is needed to clarify whether there is an impairment in OCD in the ability to disengage the DMN when transitioning from rest to task, and whether this may present a transdiagnostic trait across psychiatric disorders.

The absence of expected associations between certain clinical features of OCD and task-related activation may stem from the limited phenotypic detail of the included samples, such as past medication use, comorbidities, and symptom subtypes. This constraint prevented more sensitive subgroup analyses that might have identified important moderators of task effects.

Like all retrospective analyses of legacy data, our study was limited by the lack of harmonized data collection and reporting. Nonetheless, our results proved robust to sample effects, as leave-one-sample-out analyses did not suggest undue influence of larger samples, unlike what was previously observed in our negative valence domain analyses(12). While sensitivity analyses using frequentist models yielded less pronounced effects, Bayesian multilevel models provided more robust and reliable evidence for differences in activation by leveraging partial pooling across brain regions and eliminating the need for multiple testing corrections. Though different task paradigms captured various aspects of executive function, we demonstrated a strong executive control effect in nearly all predefined ROIs across tasks. Moreover, results were very robust to task effects, as results of the analyses on the TOL tasks alone closely matched those from the combined-task analyses, particularly at the whole-brain level. This suggests that the TOL task effectively captures the broader construct of executive function, which we were able to examine independently of inhibitory control– a limitation of all previous meta-analyses(8–10) that may explain discordant findings.

## Conclusion

Our findings support the hypothesis that the RDoC’s cognitive systems domain is relevant to OCD and that impaired goal-directed behavior in OCD may stem from insufficient activation of dorsal cognitive and frontoparietal circuits during executive control. This insufficient activation of key task-positive regions may be related to a decoupling failure of the default mode network during task performance, pointing to potential neurocircuit-level dysfunctions in OCD.

## Acknowledgments

The ENIGMA-Obsessive Compulsive Disorder Working-Group gratefully acknowledges support from the International Obsessive-Compulsive Disorder Foundation with the Innovator Award of 2021 awarded to OAvdH and CV. The authors were further supported by the Spanish Ministry of Science, Innovation and Universities (Grant No. ISCIII PI22/00752 to PA); the German Research Foundation (Deutsche Forschungsgemeinschaft); the Basic Research Program of the Korea Brain Research Institute and the National Research Foundation of Korea of the Korean Ministry of Science & ICT through the Bio & Medical Technology Development Program (Grant Nos. 2020M3E5D9079910 and 2021M3A9E408078412 to JSK) and the Brain Science Convergence Research Program (Grant Nos. RS-2023-00266120 and 21-BR-03-01 to MK); the National Research Foundation of South Africa; the South African Medical Research Council (SAMRC); the Carlos III Health Institute (Grant No. PI19/01179 to CSM); the Catalan Audiovisual Media Corporation La Fundació Marató de TV3 (Grant No. 202201 to CSM); the National Institute of Health (Grant No. R01MH138569 to PMT, OvdH, HBS); the Western Norway Health Authority (Grants Nos. 911754 and 911880 to ALT); and the Wellcome Trust/DBT India Alliance (Grant No. 500236/Z/11/Z to GV).

## Data availability

Processed and de-identified tabulated ROI-based and whole-brain parcellated brain activation data will be made available through the ENIGMA Toolbox (<u>enigma-toolbox.readthedocs.io</u>) in accordance with ENIGMA consortium policies. The code used for analyses is available at <u>github.com/nadza-dz/task-based-fMRI-processing-pipeline-ENIGMA-OCD.git</u>.

