## Supplement for "Executive control in Obsessive-Compulsive Disorder: A worldwide mega-analysis of task-based functional neuroimaging data of the ENIGMA-OCD consortium"

### Supplemental Methods

MRI acquisition parameters ……………………………………………….. 2
Contrast vectors ………………………………………………………….. 4
Regions-of-interest coordinates ……………………………………… 6
Preprocessing and quality control guidelines……………………... 7
Data exclusions ………………………………………………………….. 7
Explanation of RBA ridge plots ………………………………………. 8

### Supplemental Results

Demographics per site ………………………………………………. 9
Main effect of task load ……………………………………………... 10
Case-control effect of task load ……………………………………. 12
Clinical covariate effects of task load ……………………………... 13
Behavioral performance …………………………………………….. 14
Frequentist statistics ………………………………………………… 15
Whole-brain Bayesian multilevel results …………………………... 16
Sensitivity leave-one-sample-out analyses results ……………... 17
Sensitivity sample and task correction methods ………………... 22
Post-hoc analysis …………………………………………………….. 23

Supplemental references …………………………………………….. 24

### Supplemental Methods

**Table S1 –** Scan acquisition parameters of studies included in mega-analysis

| Site-PI | Scanner | Head coil (# channels) | Pulse sequence | Single-/ multiband | Matrix | Volumes (N) | Slices (N) | Scan order | Slice gap (mm) | TR (ms) | TE (ms) | Flip angle (°) | FOV (mm) | Voxel size (mm) | Slice time correction applied | Prospective motion correction during scanning |
|---|---|---|---|---|---|---|---|---|---|---|---|---|---|---|---|---|
| **van den Heuvel** (1) | 1.5T Siemens Sonata | standard circularly polarized | SS-EPI | singleband | 64x64 | 433 | 35 | interleaved | 0.5 | 3045 | 45 | 90 | 192 | 3x3x3 | Yes | No |
| **van den Heuvel** (2) | 3T GE Discovery MR750 | 32 | GE-EPI | singleband | 64x64 | 307 | 42 | ascending | 0.33 | 2200 | 28 | 90 | 211 | 3.3x3.3x 3 | Yes | No |
| **van den Heuvel** (3) | 3T GE Signa HDxt | 8 | GE-EPI | singleband | 64x64 | variable | 40 | ascending | 0.2 | 2100 | 30 | 80 | 240 | 3x3x3 | Yes | No |
| **Huyser** (4) | 3T Philips Intera MR | 6 | GE-EPI | singleband | 96x96 | 440 | 40 | interleaved | 0 | 2300 | 30 | 90 | 220x 120x 220 | 2.29x 2.29x3 | No | No |
| **Reddy** | 3T Seimens Skyra | 32 | SS-EPI | singleband | 64x64 | 168 | 37 | descending | 0.8 | 2000 | 30 | 78 | 192 | 3x3x3 | Yes | No |
| **Menchón/ Soriano-Mas** (5) | 1.5T GE Signa Excite | 8 | GE-EPI | singleband | 64x64 | 180 | 22 | interleaved | 1 | 2000 | 50 | 90 | 240 | 3.75x 3.75x4 | Yes | No |
| **Menchón/ Soriano-Mas** | 3T Philips Ingenia | 32 | SS-EPI | singleband | 80x80 | 97 | 40 | interleaved | 0 | 2000 | 25 | 90 | 240 | 3x3x3 | Yes | No |
| **Lazaro** | 3T Siemens Magnetom Tim Trio | 8 | GE-EPI | singleband | 80x80 | 220 | 32 | interleaved | 0 | 2000 | 29 | 80 | 240 | 3x3x4 | Yes | No |
| **Thorsen** | 3T GE Discovery MR750 | 8 | GE-EPI | singleband | 64x64 | 430 | 34 | interleaved | 0.2 | 2.1 | 30 | 80 | 220 | 3.44x 3.44x3 | Yes | No |
| **Beucke/ Kathmann** (6) | 3T Siemens Magneton Trio Tim MR | 32 | GE-EPI | singleband | 64x64 | 262 | 32 | descending | 25% | 2000 | 30 | 78 | 192 | 3.0x3.0x 3.75 | Yes | No |

| | | | | | | | | | | | | | | | |
|---|---|---|---|---|---|---|---|---|---|---|---|---|---|---|---|
| **Stein/ Lochner** | 3T Siemens Magnetom Skyra | 32 | Ep2D | singleband | 64x64 | 139 | 36 | interleaved | 0.05 | 3000 | 25 | 90 | 200 | 3.1x3.1x 3.9 | Yes | No |
| **Benedetti** | 3T Philips Gyroscan Intera | 32 | GE-EPI | singleband | 96x96 | 200 | 40 | interleaved | 0 | 3000 | 30 | 85 | 240 | 3x3x3 | Yes | No |
| **Kwon** (7) | 3T Siemens Trio | 12 | GE-EPI | singleband | 64x64 | 1080 | 27 | interleaved | 0.7 | 2000 | 30 | 90 | 240 | 3.4x3.4x 4 | No | Yes |
| **Voon** | 3T Siemens Prisma | 64 | Ep2D | singleband | 96x96 | variable (max 270) | 33 | interleaved | 0.75 | 2000 | 30 | 90 | 192 | 2x2x3 | Yes | No |
| **Stewart** (8) | 3T GE Discovery MR750 | 12 | GE-EPI | singleband | 96x96 | 412 | 41-43 | interleaved | 1 | 2000 | 25 | 90 | 256 | 3x3x3 | No | No |

**Table S2 –** Contrast vectors of executive control and task load contrasts per task

| Site-PI | City, Country, Sample | Task | Executive control contrast | Task load contrast* |
|---|---|---|---|---|
| van den Heuvel | Amsterdam, NDL I | Tower of London | 5-step [1]<br>4-step [1]<br>3-step [1]<br>2-step [1]<br>1-step [1]<br>Count [-5] | 5-step [2]<br>4-step [1]<br>3-step [0]<br>2-step [-1]<br>1-step [-2]<br>Count [0] |
| van den Heuvel | Amsterdam, NDL II | N-Back | 3-back [1]<br>2-back [1]<br>1-back [1]<br>0-back [-3] | 3-back [1]<br>2-back [0]<br>1-back [-1]<br>0-back [0] |
| van den Heuvel | Amsterdam, NDL III | Tower of London | 5-step [1]<br>4-step [1]<br>3-step [1]<br>2-step [1]<br>1-step [1]<br>Count [-5] | 5-step [2]<br>4-step [1]<br>3-step [0]<br>2-step [-1]<br>1-step [-2]<br>Count [0] |
| Huyser | Amsterdam, NDL IV | Tower of London | 5-step [1]<br>4-step [1]<br>3-step [1]<br>2-step [1]<br>1-step [1]<br>Count [-5] | 5-step [2]<br>4-step [1]<br>3-step [0]<br>2-step [-1]<br>1-step [-2]<br>Count [0] |
| Reddy | Bangalore, IND | N-Back | 2-back [1]<br>1-back [1]<br>0-back [-2] | — |
| Menchón/Soriano-Mas | Barcelona, ESP I | Task-switching | Task switch [1]<br>Task repeat [-1] | — |
| Menchón/Soriano-Mas | Barcelona, ESP II | Task-switching | Task switch [1]<br>Task repeat [-1] | — |
| Lazaro | Barcelona, ESP III | Task-switching | Task switch [1]<br>Task repeat [-1] | — |
| Thorsen | Bergen, NOR | Tower of London | 5-step [1]<br>4-step [1]<br>3-step [1]<br>2-step [1]<br>1-step [1]<br>Count [-5] | 5-step [2]<br>4-step [1]<br>3-step [0]<br>2-step [-1]<br>1-step [-2]<br>Count [0] |
| Beucke/Kathmann | Berlin, GER | N-Back | 3-back [1]<br>2-back [1]<br>1-back [1]<br>0-back [-3] | 3-back [1]<br>2-back [ 0]<br>1-back [-1]<br>0-back [0] |
| Stein/Lochner | Cape Town, ZAF | N-Back | 1-back [1]<br>0-back [-1] | — |

| | | | | |
|---|---|---|---|---|
| Benedetti | Milano, ITA | Tower of London | 5-step [1]<br>4-step [1]<br>3-step [1]<br>2-step [1]<br>1-step [1]<br>Count [-5] | — |
| Kwon | Seoul, KOR | Tower of London | 4-step [1]<br>3-step [1]<br>2-step [1]<br>1-step [1]<br>Count [-4] | 4-step [2]<br>3-step [1]<br>2-step [-1]<br>1-step [-2]<br>Count [0] |
| Voon | Shanghai, CHN | N-Back | 2-back [1]<br>1-back [1]<br>0-back [-2] | 2-back [ 1]<br>1-back [-1]<br>0-back [0] |
| Stewart | Vancouver, CAN | Tower of London | 5-step [1]<br>4-step [1]<br>3-step [1]<br>2-step [1]<br>1-step [1]<br>Count [-5] | 5-step [2]<br>4-step [1]<br>3-step [0]<br>2-step [-1]<br>1-step [-2]<br>Count [0] |

* For tasks in which there was an odd number of task load conditions, the middle condition was given a weight of 0 in the task load contrast. This was done to create equally spaced steps along the task load levels, enabling consistent contrasts of high versus low load across task paradigms. This meant that the middle task load condition and control trials together formed the implicit baseline in these tasks.

**Table S3 -** Regions of interest identified from Nitschke et al. (2017; (9)) showing whole-brain activation in healthy controls during the Tower of London task for the planning and task load contrasts. Coordinates are in MNI152 NLIN 6th generation space. dlPFC = dorsolateral prefrontal cortex; FEF = frontal eye fields; SMA = supplementary motor area; IPL = inferior parietal lobule; rlPFC = rostrolateral prefrontal cortex

| Region | Lateralization | PLANNING | | | TASK LOAD | | |
|---|---|---|---|---|---|---|---|
| | | x | y | z | x | y | z |
| dlPFC | Left | −40.6 | 31.8 | 30.6 | −43.2 | 32.8 | 30.9 |
| | Right 1 | 41.4 | 35.5 | 29.9 | 42.2 | 38.7 | 27.3 |
| | Right 2 | – | – | – | 46 | 26.1 | 42.8 |
| FEF | Left 1 | −26.2 | 8.7 | 57.1 | −24.8 | −0.9 | 65.2 |
| | Left 2 | – | – | – | −25.0 | 22.5 | 53.3 |
| | Right 1 | 29.3 | 11.3 | 55.2 | 29.6 | 17.5 | 55.9 |
| | Right 2 | – | – | – | 22.8 | −.5 | 66 |
| SMA/pre-SMA | Bilateral | −1.9 | 25.8 | 43.9 | −3.5 | 24.5 | 45.6 |
| IPL | Left 1 | −38.2 | −45.6 | 44.9 | −53.8 | −40.7 | 48.8 |
| | Left 2 | – | – | – | −40.5 | −51.9 | 51 |
| | Right | 46.3 | −39.8 | 47.2 | 51.2 | −42.8 | 46.7 |
| Precuneus | Bilateral | 0 | −61.5 | 57.3 | −11.1 | −57.7 | 60.9 |
| | Right 1 | 32.9 | −72.1 | 42.5 | 8.3 | −59.3 | 58.4 |
| | Right 2 | – | – | – | 42.4 | −74.8 | 39.3 |
| Caudate | Left | −12.2 | 14.3 | 1 | −16.4 | 7.5 | 11.1 |
| | Right 1 | 13.9 | 10.1 | 2.6 | 18.3 | 6.8 | 15.6 |
| | Right 2 | – | – | – | 14.1 | 5.2 | −2.1 |
| Anterior insula | Left | −31.0 | 23.8 | −0.9 | −32.9 | 23.1 | −4.5 |
| | Right | 33.1 | 24.3 | −4.2 | – | – | – |
| rlPFC | Left | −35.3 | 55.8 | 5.6 | −40.0 | 55.5 | 10.4 |
| | Right | 33.7 | 57.6 | 3.8 | – | – | – |
| Posterior cingulate | Left | −17.1 | −58.2 | 21.5 | – | – | – |
| | Right | 17.1 | −58.2 | 21.5 | – | – | – |
| Inferior Occipital Gyrus | Left | −43.3 | −79.2 | −4.1 | – | – | – |
| | Right | 43.3 | −79.2 | −4.1 | – | – | – |
| Inferior Temporal Gyrus | Left | −52.1 | −62.1 | −5.7 | – | – | – |
| | Right | 52.1 | −62.1 | −5.7 | – | – | – |

### Preprocessing and quality control guidelines

Preprocessing was done at each site according to harmonized guidelines and each site performed quality-control on their own preprocessed data following harmonized guidelines which can be retrieved from the online repository 10.5281/zenodo.14800135.

### Data exclusions

For six (6) participants (3 OCD and 3 HCs) the Independent Component Analysis (ICA-AROMA) portion of the analysis pipeline failed to run, twenty-nine (29) participants (10 OCD and 19 HCs) failed quality assessment due to poor skull-stripping or registration, and six (6) participants (3 OCD and 3 HCs) were excluded due to excessive motion.

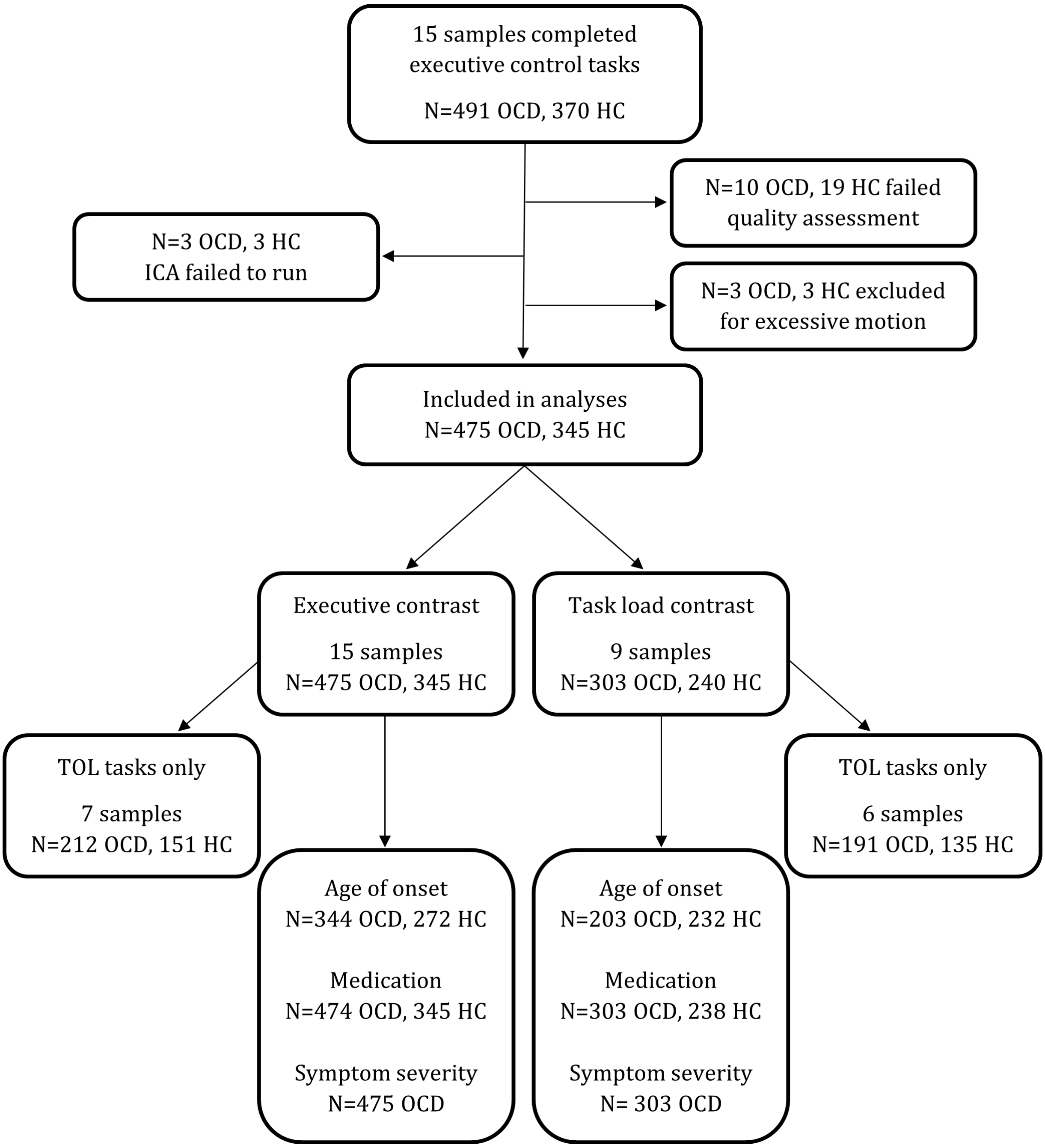

**Explanation of how to interpret ridge plot results of Regional Bayesian Analysis**

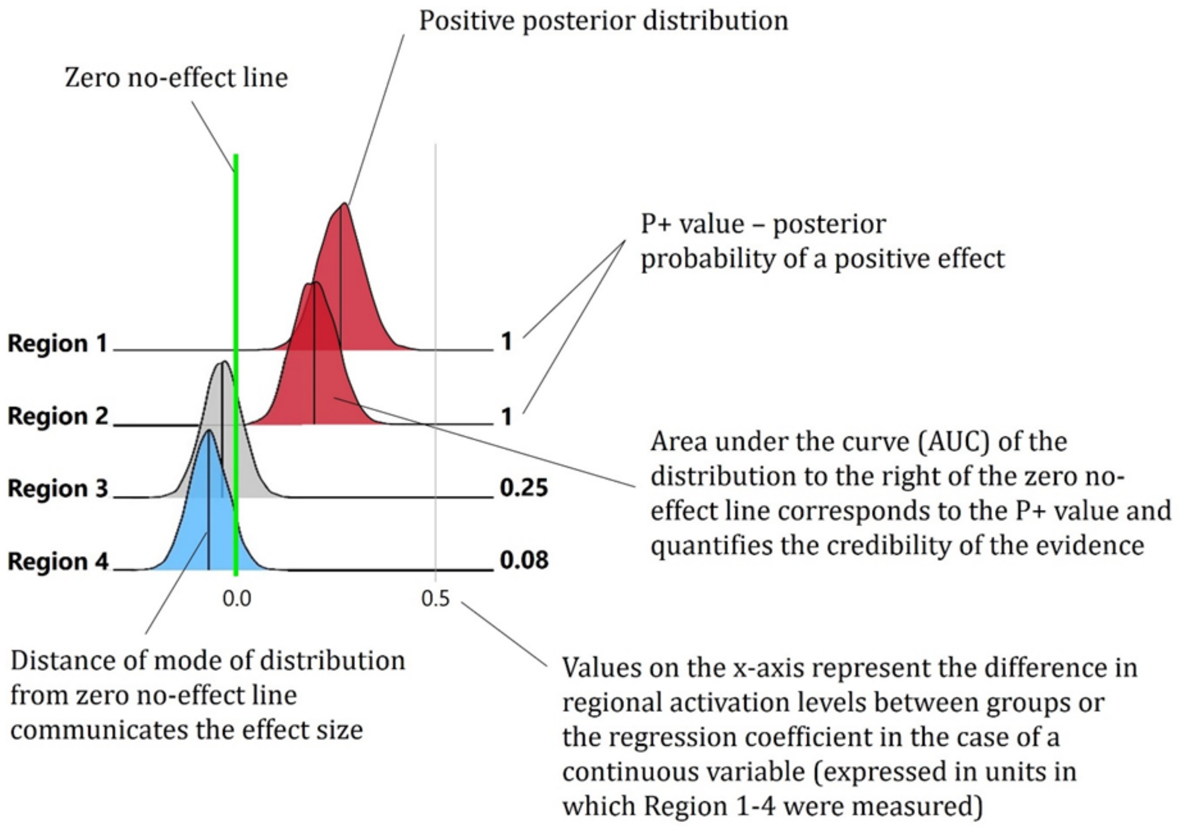


Consider this hypothetical analysis of effects in four regions of interest as performed under Regional Bayesian Analysis (RBA; Chen et al., 2019 (10)). In Regions 1 and 2 there is very strong evidence of a positive effect as the entire AUC is to the right of the no-effect line (P+=1.0), denoting a nearly 100% probability of the effect being positive in these regions. Region 1 has a larger effect size than Region 2 as the mode of the distribution is further away from the no-effect line. In Region 4 there is moderate evidence for a negative effect as AUC is largely to the left of the no-effect line (P+=0.08). Small values of P+ convey evidence that the effect is negative – a P+ value of 0.08 indicates that the probability of the effect being positive is only 8%, so the probability of it being negative is 92%. For Region 3, the distribution crosses the no-effect line with 25% of the AUC to the right of the no effect line (P+=0.25), meaning there is no strong evidence of a positive effect, but also no strong evidence of a negative effect in this region

### Supplemental Results

**Table S4 - Demographics per site**

| Site-PI | City, Country, Sample | n OCD | n HC | % Male | Age (mean ± SD) | % Child onset OCD | % Medicated OCD | (C)Y-BOCS (mean ± SD) | TOL accuracy (mean ± SD) |
|---|---|---|---|---|---|---|---|---|---|
| van den Heuvel | Amsterdam, NDL I | 20 | 20 | 37.5 | 31.62 ± 8.34 | 50 | 0 | 22.26 ± 6.76 | 89.92 ± 9.75 |
| van den Heuvel | Amsterdam, NDL II | 39 | 33 | 51.39 | 38.86 ± 10.27 | 61.54 | 0 | 21 ± 6.07 | — |
| van den Heuvel | Amsterdam, NDL III | 65 | 25 | 37.78 | 36.43 ± 12.41 | 55.38 | 64.62 | 28.66 ± 4.63 | 90.61 ± 6.46 |
| Huyser | Amsterdam, NDL IV | 27 | 23 | 38 | 13.7 ± 2.38 | 96.3 | 0 | 24.58 ± 5.07 | 90.15 ± 5.8 |
| Reddy | Bangalore, IND | 38 | 27 | 61.54 | 26.89 ± 5.5 | 42.11 | 31.58 | 24.24 ± 7.87 | — |
| Menchón/Soriano-Mas | Barcelona, ESP I | 18 | 18 | 52.78 | 46.06 ± 9.07 | 50 | 94.44 | 23.28 ± 5.84 | — |
| Menchón/Soriano-Mas | Barcelona, ESP II | 21 | 0 | 42.86 | 35.29 ± 10.08 | 57.14 | 100 | 24.52 ± 6.42 | — |
| Lazaro | Barcelona, ESP III | 60 | 35 | 55.79 | 15.28 ± 2.14 | 100 | 80 | 19 ± 7.43 | — |
| Thorsen | Bergen, NOR | 38 | 28 | 37.88 | 30.27 ± 9.49 | 39.47 | 23.68 | 26.32 ± 4.33 | 87.09 ± 7.75 |
| Beucke/Kathmann | Berlin, GER | 49 | 45 | 46.81 | 31.86 ± 8.66 | 65.31 | 42.86 | 23.08 ± 5.24 | — |
| Stein/Lochner | Cape Town, ZAF | 14 | 9 | 34.78 | 32.3 ± 11.73 | 64.29 | 85.71 | 21.93 ± 5.31 | — |
| Benedetti | Milano, ITA | 21 | 15 | 63.89 | 34.36 ± 10.59 | 109.52 | 76.19 | 31.45 ± 4.38 | 59.38 ± 18.54 |
| Kwon | Seoul, KOR | 17 | 21 | 60.53 | 26.18 ± 5.54 | 70.59 | 0 | 30.35 ± 4.14 | 83.57 ± 12.34 |
| Voon | Shanghai, CHN | 24 | 27 | 58.82 | 32.49 ± 8.22 | 50 | 70.83 | 23.79 ± 7.4 | — |
| Stewart | Vancouver, CAN | 24 | 19 | 37.21 | 14.34 ± 3.23 | 100 | 79.17 | 13.42 ± 7.86 | 91.5 ± 8.49 |

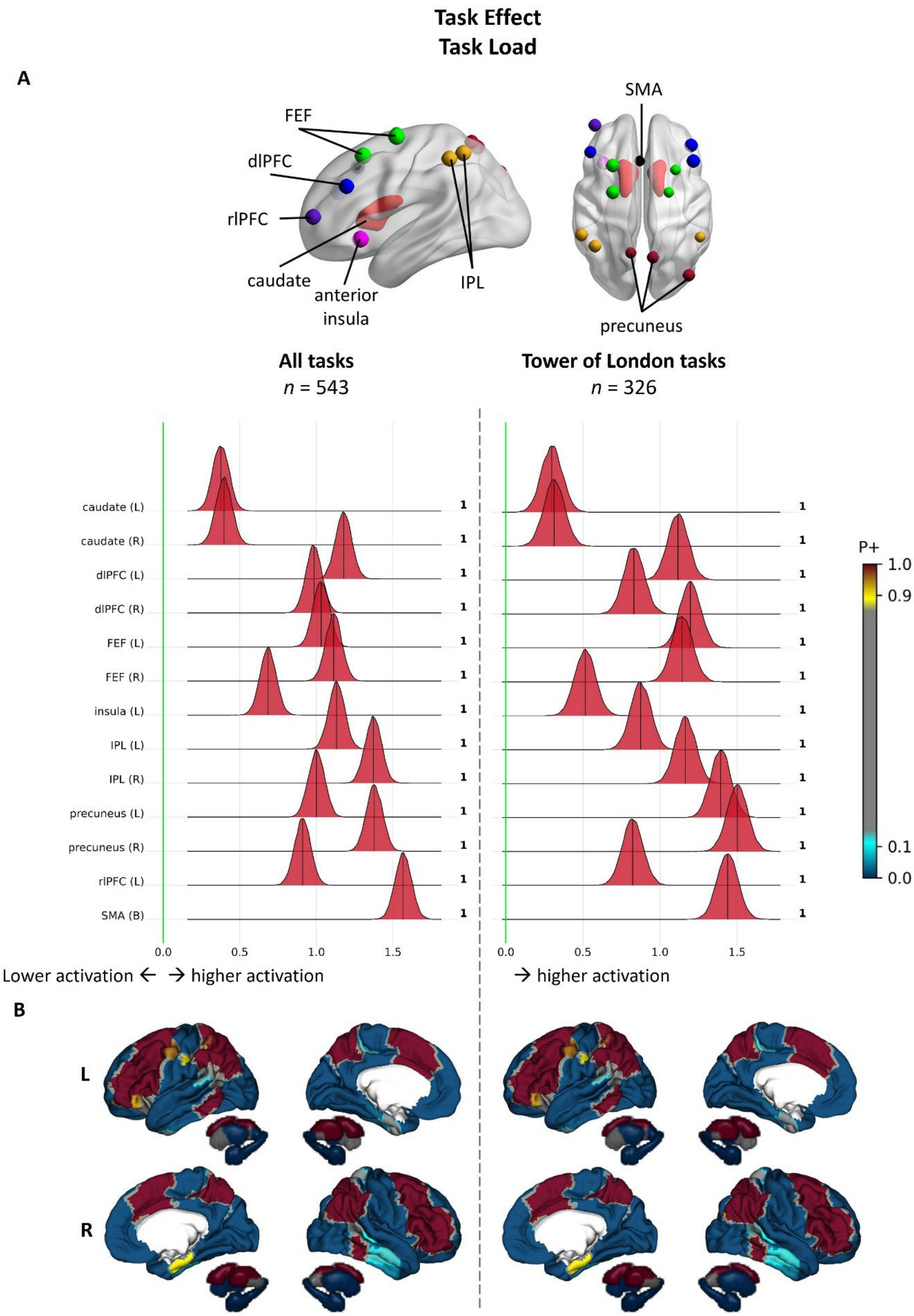


***Figure S1 - Main effect of task load in executive control in all participants***(A) Region-of-interest results from Bayesian multilevel analyses of group-level executive contrast across individuals with OCD and HCs in all tasks (N tasks = 9, n participants = 543) and in only Tower of London tasks (N tasks = 6, n participants

= 326). Posterior probability distributions express the credibility of an effect in each region. Next to each distribution the posterior probability of a positive effect (P+) is shown. In regions with posterior distributions to the right of the green no-effect line there is evidence of stronger activation during executive control while in regions with posterior distributions to the left of this line there is evidence of weaker activation during executive control. Regions are color-coded to reflect the strength of evidence for an effect, where in (darker) red regions there is stronger evidence of activation during executive control (P+ values >0.90 indicate moderate to very strong evidence for a positive effect). In (darker) blue regions there is stronger evidence of deactivation during executive control (P+ values <0.10 indicate moderate to very strong evidence for a negative effect). In grey regions there is no strong evidence of activation or deactivation during executive control. Values on the x-axis represent the difference in regional activation levels between executive and control trials (expressed as difference in Z-scores). (B) Whole-brain effects from Bayesian multilevel analyses. P+ values derived from Bayesian multilevel analyses denote the probability that there is stronger brain activation in a given region of the Schaefer 200-parcel 7-network cortical atlas and Melbourne 32-region subcortical atlas. Displayed are lateral and medial views of the cortex. dlPFC= dorsolateral prefrontal cortex; FEF = frontal eye fields; IOG = inferior anterior occipital gyrus; IPL = inferior parietal lobule; ITG = inferior temporal gyrus; rlPFC = rostrolateral prefrontal cortex; SMA = supplementary motor area.

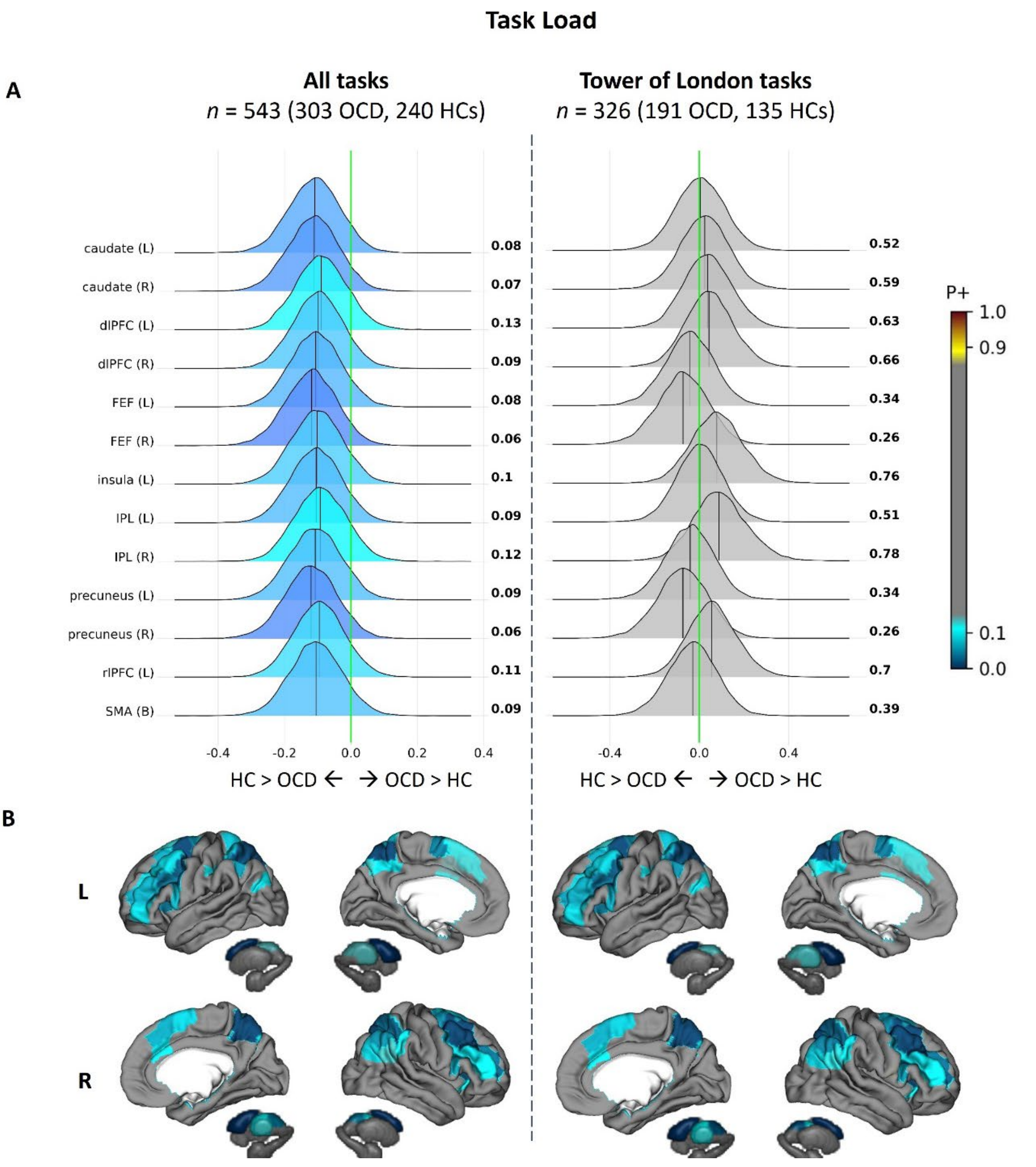


***Figure S2 - Case-control differences in task load in executive control*** (A) Region-of-interest results from Bayesian multilevel analyses. Distributions to the right of the green no-effect line represent regions in which individuals with OCD show evidence for stronger activation than HCs. (Darker) red color represents regions in which individuals with OCD show moderate-to-very-strong evidence for stronger activation than HCs. Regions with posterior distributions to the left of the no-effect line show evidence for stronger activity in HCs than in OCD. (Darker) blue color represents regions in which HCs show moderate-to-very-strong evidence for stronger activation than OCD individuals. In gray-colored regions there is no evidence of a difference between HCs and OCD individuals. Values on the x-axis represent the difference in regional activation levels between HCs and OCD (expressed as difference in Z-scores). (B) Whole-brain effects from Bayesian multilevel analyses. dlPFC= dorsolateral prefrontal cortex; FEF = frontal eye fields; IOG = inferior anterior occipital gyrus; IPL = inferior parietal lobule; ITG = inferior temporal gyrus; rlPFC = rostrolateral prefrontal cortex; SMA = supplementary motor area.

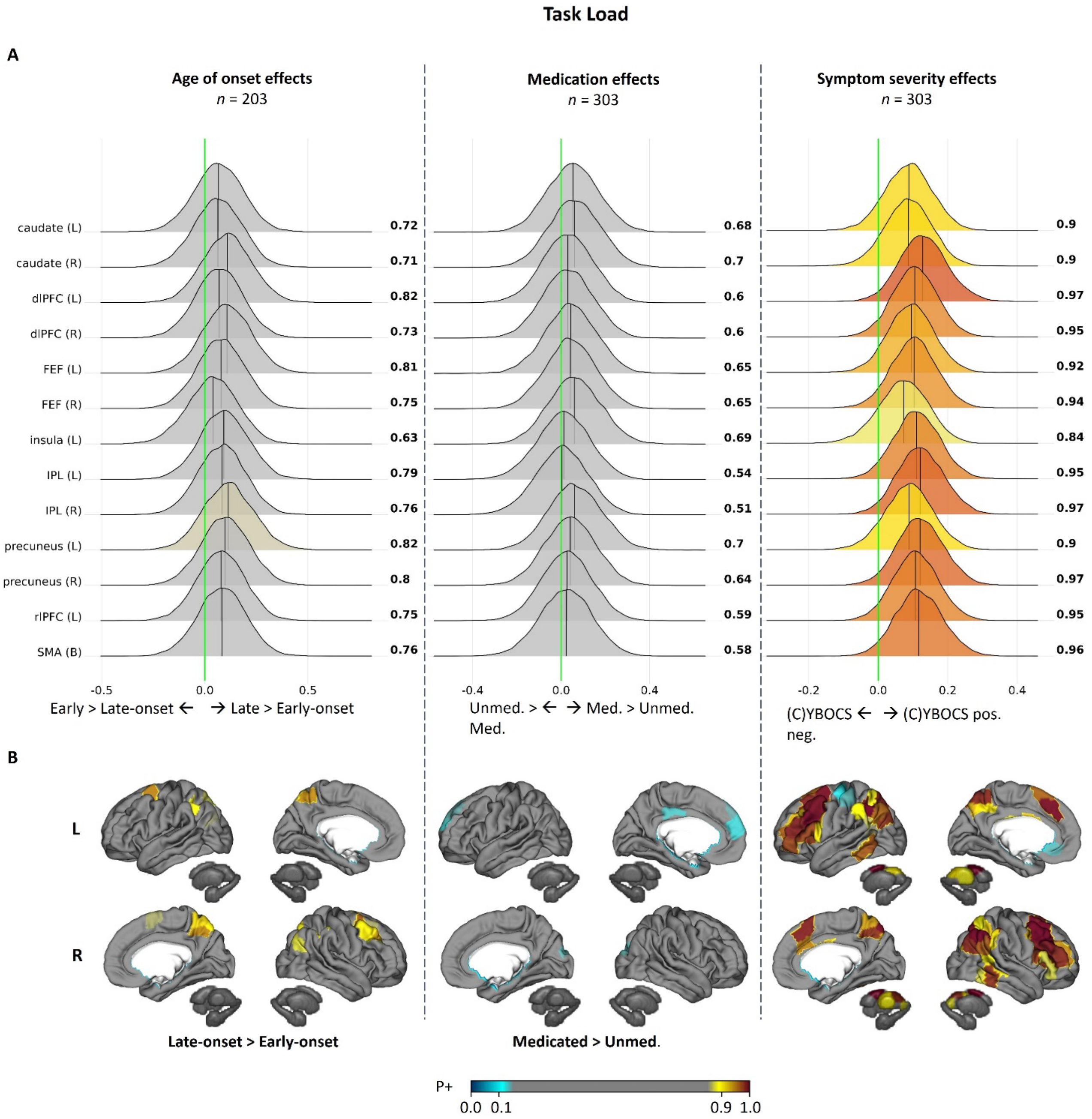


***Figure S3 - Effects of clinical features of OCD on task load in executive control***

(A) Region-of-interest effects of age of onset, medication status, and symptom severity during executive control. (B) Whole-brain analyses of executive control. (C)YBOCS = (Children's) Yale-Brown Obsessive-Compulsive Scale; dlPFC= dorsolateral prefrontal cortex; FEF = frontal eye fields; IOG = inferior anterior occipital gyrus; IPL = inferior parietal lobule; ITG = inferior temporal gyrus; rlPFC = rostrolateral prefrontal cortex; SMA = supplementary motor area.

### Behavioral performance

We only investigated behavioral performance in the Tower of London (TOL) tasks because other tasks (n-back and task-switching) controlled performance by modulating task difficulty or had too much variability in design across samples to allow a clear comparison. Though TOL tasks differed in how many steps were required to solve the most difficult trials, we calculated average accuracy per participant across all trials, and compared performance between individuals with OCD and healthy controls via t-tests. Across samples we found no evidence for difference in TOL accuracy between individuals with OCD and HCs, except in one sample (Seoul, KOR; $T_{(36)}$=2.01, $p$=0.05) (Table S2).

**Table S5 – Behavioral performance metrics of samples in which participants performed a Tower of London task**

| Site-PI | City, Country, Sample | OCD Accuracy (mean ± SD) | HC Accuracy (mean ± SD) | Statistic |
|---|---|---|---|---|
| van den Heuvel | Amsterdam, NDL I | 87.9 ± 10.04 | 91.94 ± 9.26 | $T_{(38)}$ = 1.32, P = 0.19 |
| van den Heuvel | Amsterdam, NDL III | 90.11 ± 7.08 | 91.91 ± 4.3 | $T_{(88)}$ = 1.18, P = 0.24 |
| Huyser | Amsterdam, NDL IV | 90.53 ± 5.6 | 89.71 ± 6.12 | $T_{(48)}$ = -0.49, P = 0.63 |
| Thorsen | Bergen, NOR | 87.38 ± 8.06 | 86.69 ± 7.43 | $T_{(64)}$ = -0.36, P = 0.72 |
| Benedetti | Milano, ITA | 55.75 ± 17 | 64.44 ± 19.97 | $T_{(34)}$ = 1.41, P = 0.17 |
| Kwon | Seoul, KOR | 79.27 ± 15.25 | 87.06 ± 8.21 | $T_{(36)}$ = 2.01, P = 0.05 * |
| Stewart | Vancouver, CAN | 90.58 ± 8.76 | 92.66 ± 8.21 | $T_{(41)}$ = 0.79, P = 0.43 |

* = statistically significant at an alpha level of 0.05

### Frequentist statistics

When estimating executive control-related differences in activation between individuals with OCD and HCs using a frequentist multilevel mixed effects model with a random intercept for sample and subject (rather than a Bayesian multilevel model), individuals with OCD had significantly weaker activation than HCs in the right frontal eye fields ($Z_{(16,360)}$ =-1.975, $p$=0.048), left inferior parietal lobule ($Z_{(16,360)}$ = -2.073, $p$=0.038), right inferior parietal lobule ($Z_{(16,360)}$ = -2.939, $p$=0.003), and bilateral precuneus ($Z_{(16,360)}$ = -2.066, $p$=0.039).

**Table S6 – Results of frequentist multilevel analysis of activation differences in predefined ROIs between individuals with OCD and HC during executive control**

| Region | B [SE] | Z (df=16,360) | Cohen's *d* | *p* |
|---|---|---|---|---|
| caudate (L) | -0.053 [0.111] | -0.478 | -0.040 | 0.633 |
| caudate (R) | -0.112 [0.111] | -1.009 | -0.084 | 0.313 |
| cingulate (L) | 0.152 [0.111] | 1.362 | 0.113 | 0.173 |
| cingulate (R) | 0.039 [0.111] | 0.349 | 0.029 | 0.727 |
| dlPFC (L) | -0.096 [0.111] | -0.866 | -0.072 | 0.386 |
| dlPFC (R) | -0.115 [0.111] | -1.033 | -0.086 | 0.302 |
| FEF (L) | -0.148 [0.111] | -1.326 | -0.110 | 0.185 |
| FEF (R) | -0.22 [0.111] | -1.975 | -0.164 | 0.048 * |
| insula (L) | -0.016 [0.111] | -0.147 | -0.012 | 0.883 |
| insula (R) | -0.179 [0.111] | -1.606 | -0.133 | 0.108 |
| IOG (L) | 0.197 [0.111] | 1.766 | 0.147 | 0.077 |
| IOG (R) | 0.094 [0.111] | 0.840 | 0.070 | 0.401 |
| IPL (L) | -0.231 [0.111] | -2.073 | -0.172 | 0.038 * |
| IPL (R) | -0.327 [0.111] | -2.939 | -0.244 | 0.003 * |
| ITG (L) | -0.151 [0.111] | -1.358 | -0.113 | 0.175 |
| ITG (R) | -0.005 [0.111] | -0.044 | -0.004 | 0.965 |
| precuneus (B) | -0.23 [0.111] | -2.066 | -0.171 | 0.039 * |
| precuneus (R) | -0.155 [0.111] | -1.394 | -0.116 | 0.163 |
| rlPFC (L) | 0.016 [0.111] | 0.142 | 0.012 | 0.887 |
| rlPFC (R) | -0.141 [0.111] | -1.264 | -0.105 | 0.206 |
| SMA (B) | -0.159 [0.111] | -1.433 | -0.119 | 0.152 |

* = statistically significant at an alpha level of 0.05

### Figure S4 - Whole-brain Schaefer 200-parcel cortical atlas & Melbourne 32 subcortical atlas Bayesian multilevel results

The full output of the whole-brain Bayesian multilevel models presented in Figures 1B, 2B and 3B of the manuscript can be retrieved from the online repository 10.5281/zenodo.14800135.

For each of the two contrasts of interest: 1) Executive function and 2) Task load, the following models are obtained: Intercept model; Diagnosis group effect; Age of onset effect – Late-onset vs. HC – Early-onset vs. HC – Late-onset vs. Early-onset; Medication effect – Unmedicated vs. HC – Medicated vs. HC – Medicated vs. Unmedicated; OCD severity effect.

**Figure S5 - Sensitivity leave-one-sample-out analyses for Bayesian multilevel region-of-interest results**

Executive function

*Group effect*

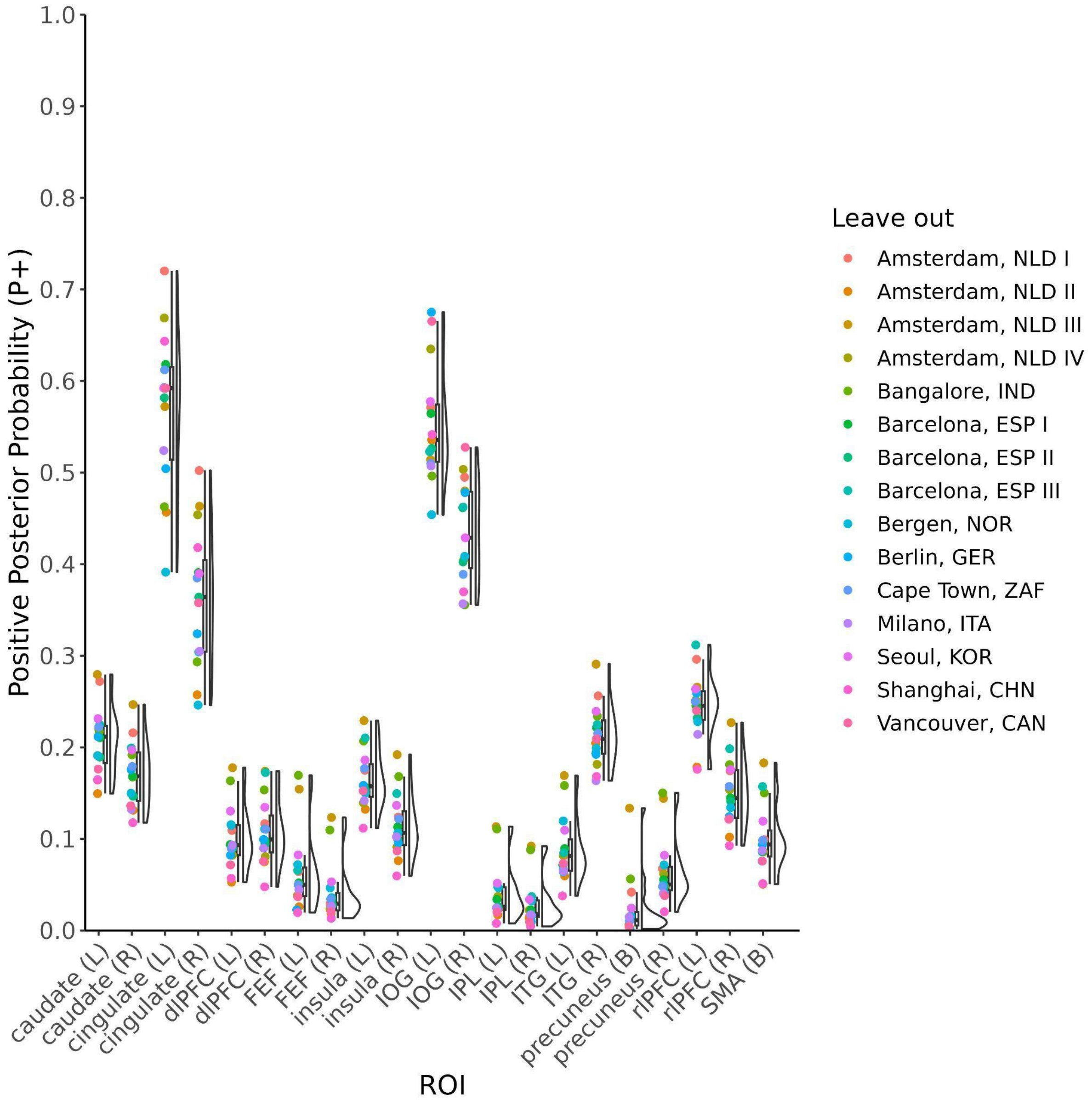

*Age of onset effect – Late-onset vs. HCs*

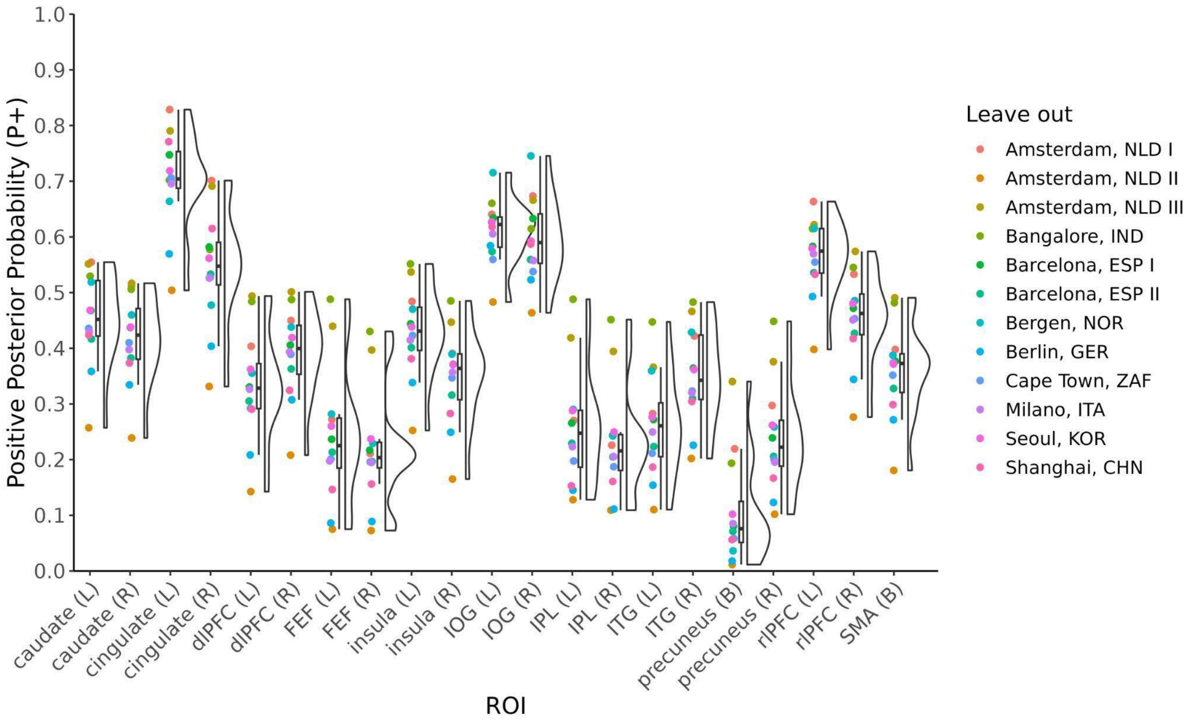


*Age of onset effect – Early-onset vs. HCs*

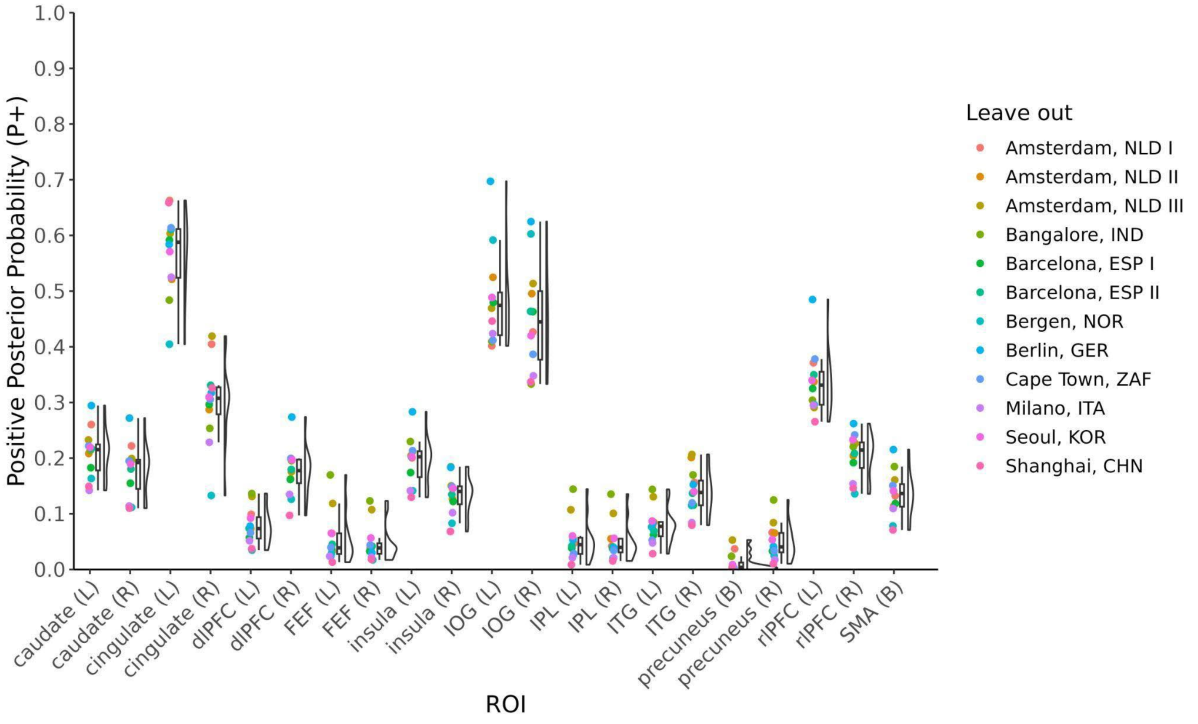

*Age of onset effect – Late-onset vs. Early-onset*

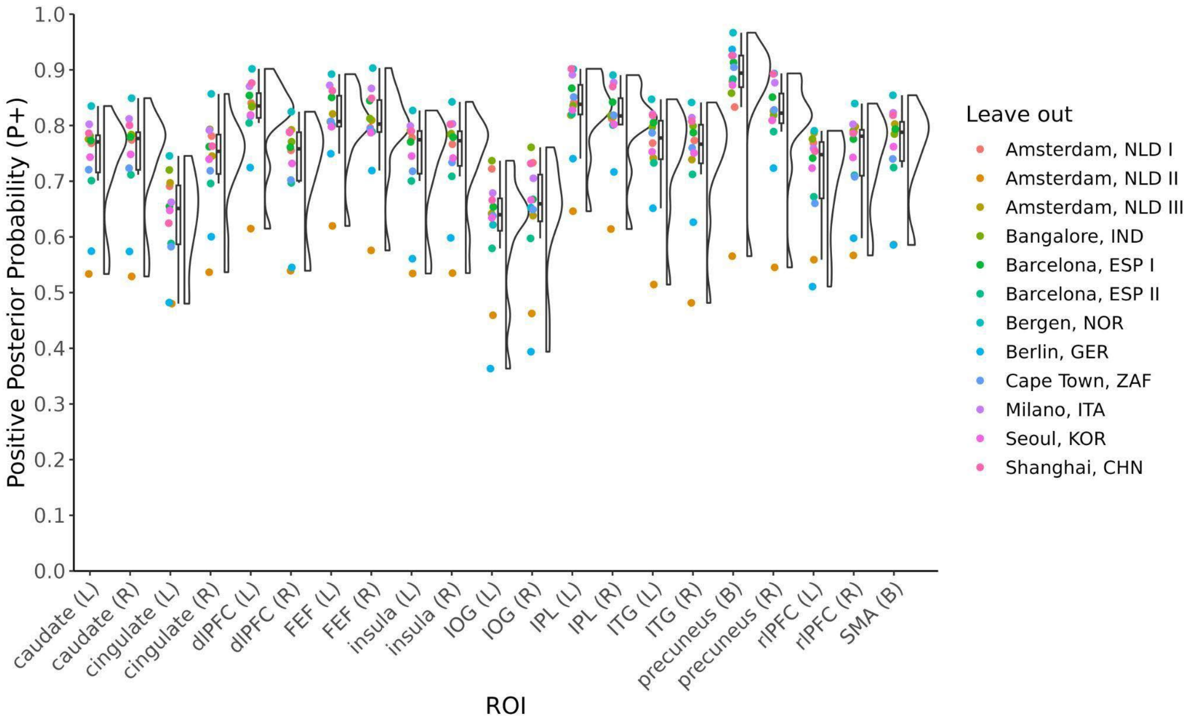


*Medication effect – Unmedicated vs. HCs*

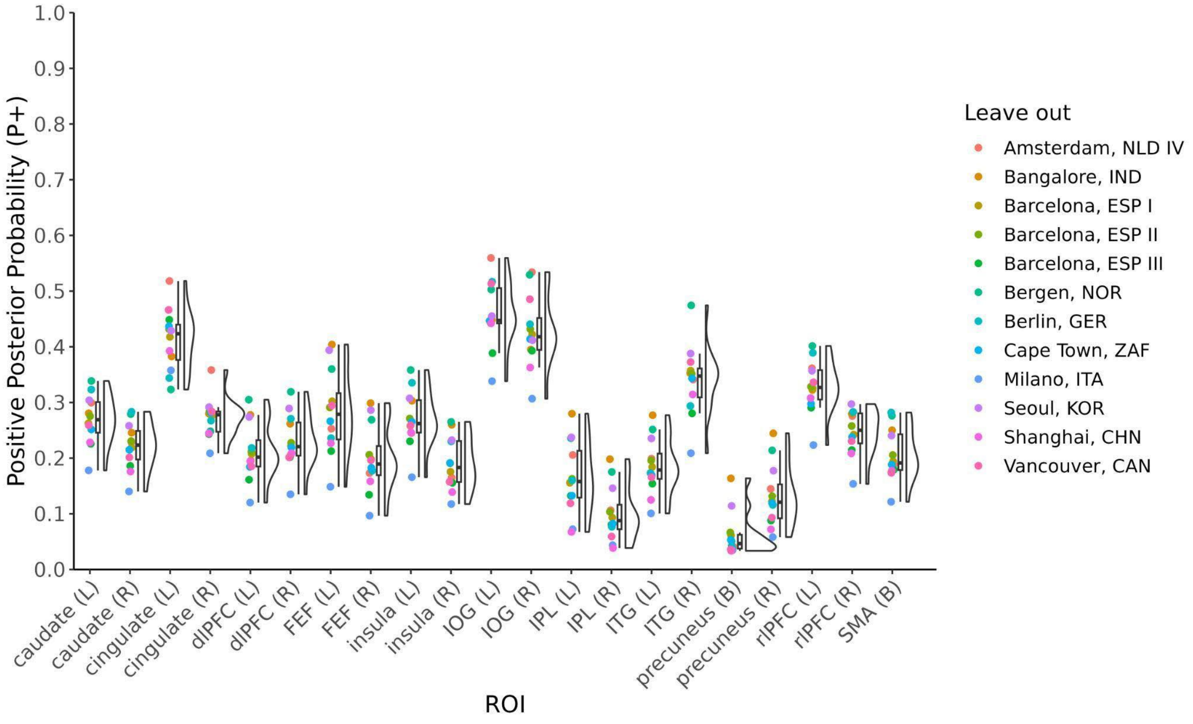

*Medication effect – Medicated vs. HCs*

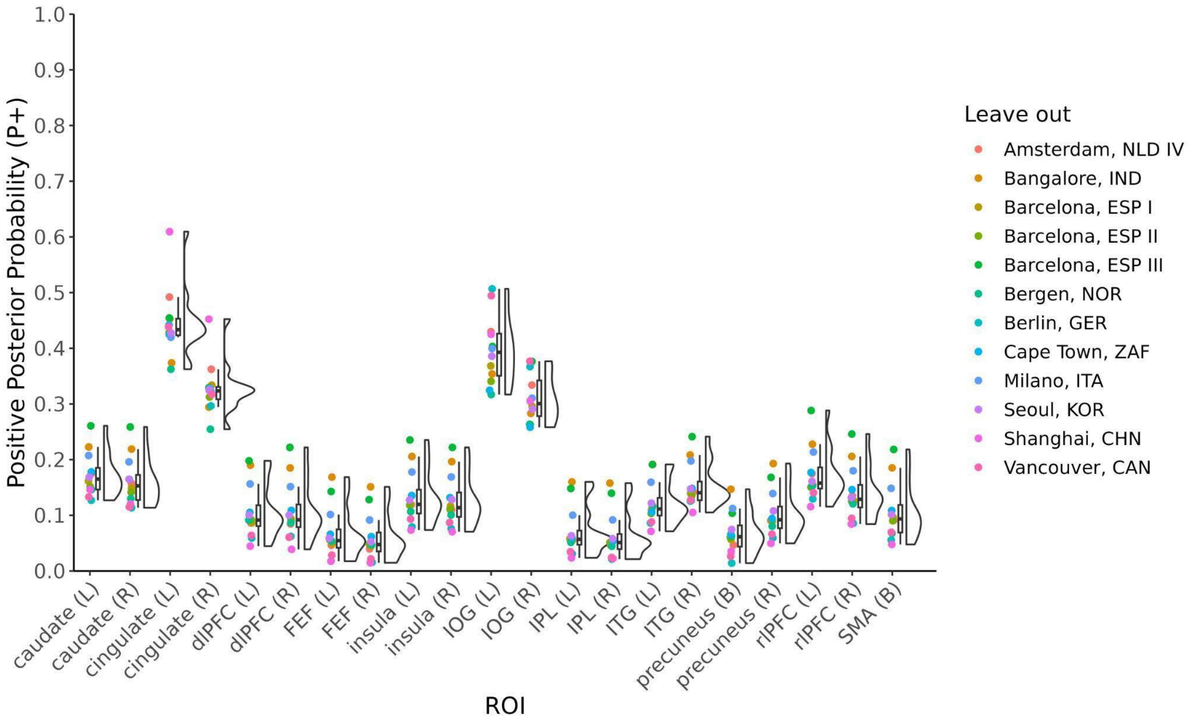


*Medication effect – Medicated vs. Unmedicated*

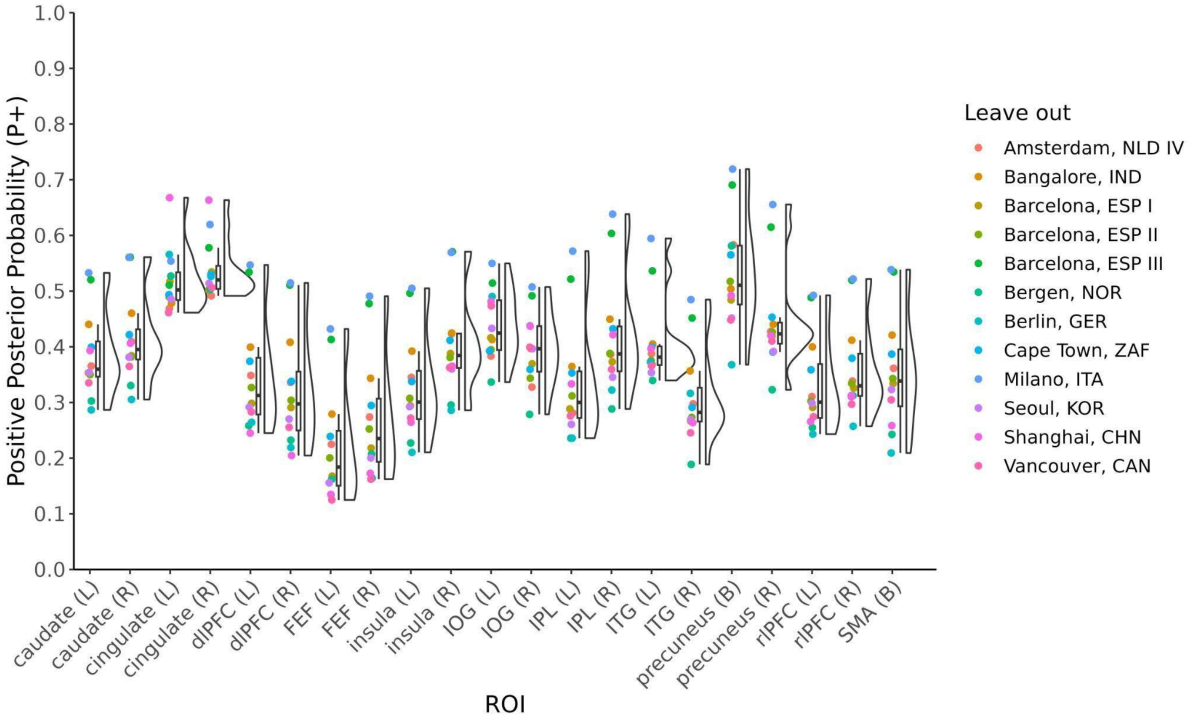

*Severity effect*

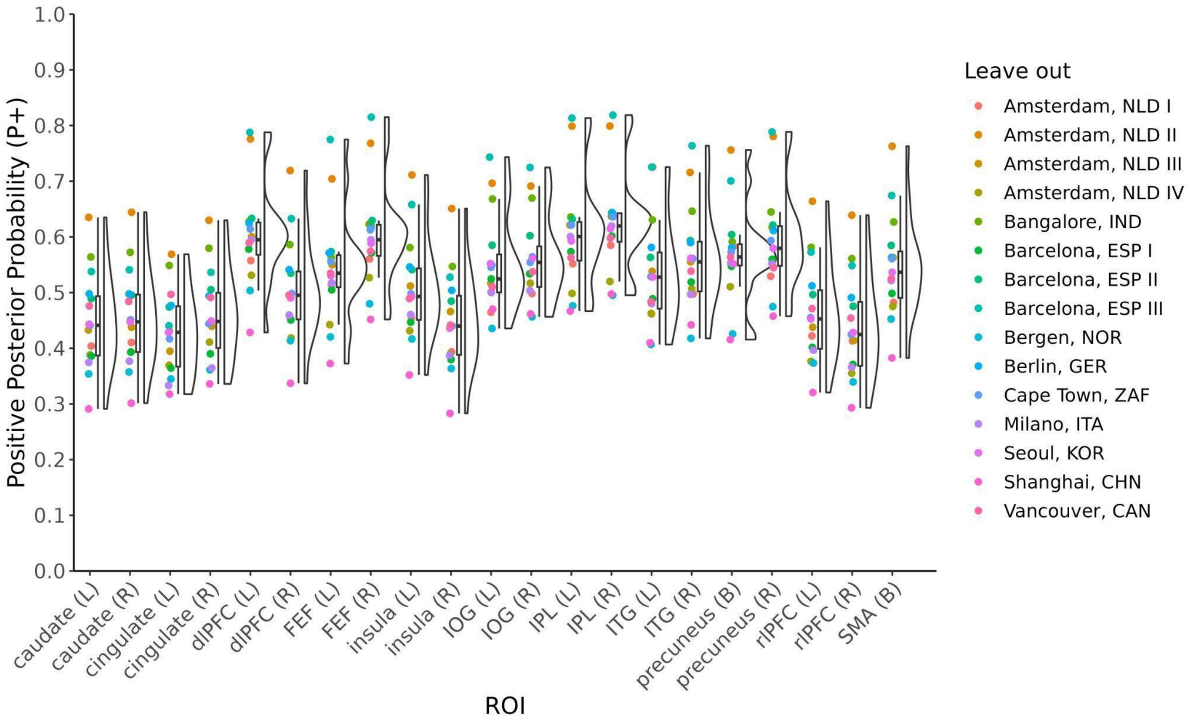

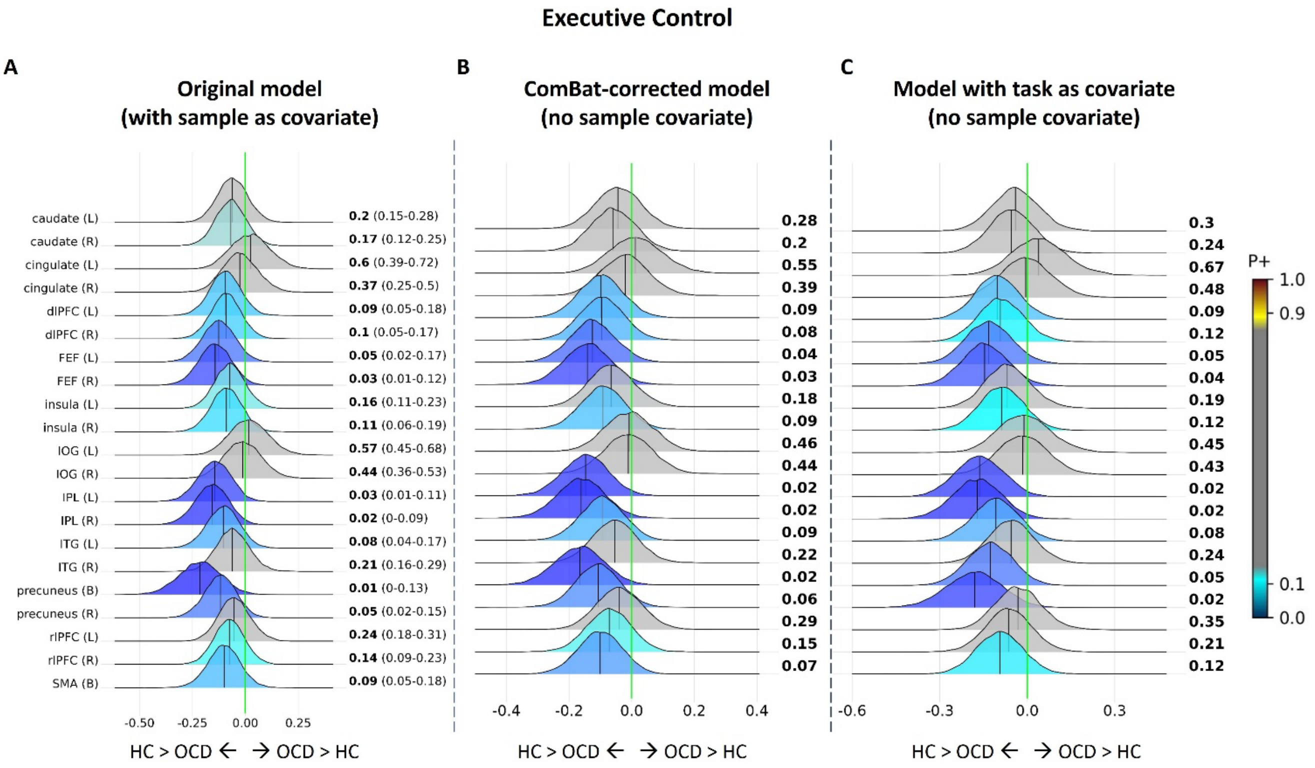


**Figure S6 - Sensitivity analyses of results across different sample and task correction methods** (A) Original region-of-interest results from Bayesian multilevel analyses of case-control effects in executive control contrast. (B) Results after ComBat harmonization to correct for sample (batch) effects, with diagnosis, age, and sex included as biological covariates in the harmonization; the Bayesian model was run on the corrected data without including sample as a covariate. (C) Results from the original dataset with task type included as a covariate in the Bayesian model instead of sample to account for task-related variance. dlPFC= dorsolateral prefrontal cortex; FEF = frontal eye fields; IOG = inferior anterior occipital gyrus; IPL = inferior parietal lobule; ITG = inferior temporal gyrus; rlPFC = rostrolateral prefrontal cortex; SMA = supplementary motor area.

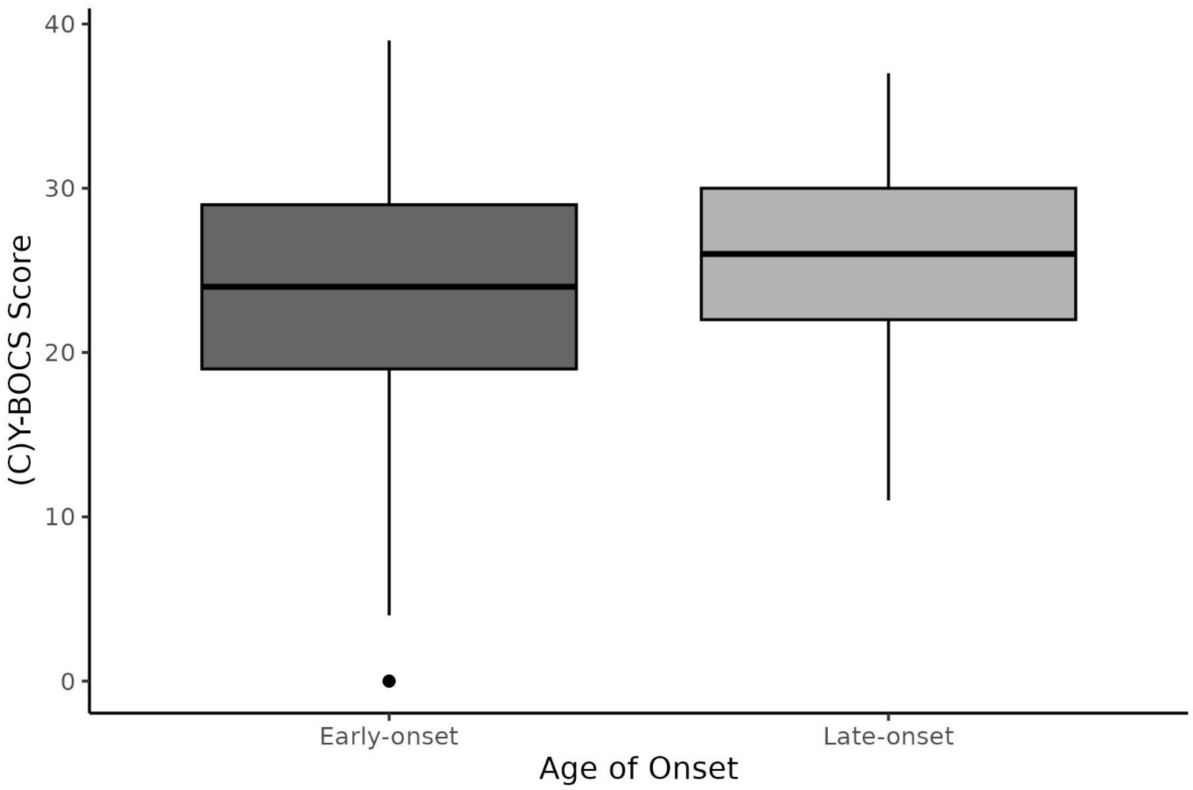


**Figure S7 - Post-hoc comparison of symptom severity scores between age of onset groups** In the subset of participants included in the task load activation analysis, we compared symptom severity ((C)Y-BOCS scores) between early- and late-onset OCD groups. Individuals with late-onset OCD showed significantly greater symptom severity than those with early-onset OCD ($T_{(277)} = 2.211$, $p = 0.028$). This group difference may underlie the positive association between symptom severity and task load activation observed in the main analysis.
